# Metabolic signals regulate resuscitation speed of antibiotic persister bacteria during infection

**DOI:** 10.64898/2026.05.05.722882

**Authors:** Jeong Moo Han, Jueun Lee, Jiseon Kim, Yi Qing Lee, Dongjoo Lee, Si Nae Lee, Duhyun Ko, Mingyeong Kang, Takyoung Yu, Jisun Lee, Kyeonghun Jeong, Eunmi Chu, Bora Shin, Lokanand Koduru, Yeji Bang, Dohyun Han, Kwangsoo Kim, Dong-Yup Lee, Ki-Wook Kim, Geum-Sook Hwang, Jinki Yeom

## Abstract

All living organisms adjust their metabolism in response to environmental changes. Under unfavorable conditions, organisms enter a state of dormancy by halting metabolism, enabling survival. Dormant bacteria become highly tolerant to antibiotics-a phenomenon called persistence. Here, we demonstrate that selective metabolic reprogramming controls the resuscitation speed of persister after antibiotic exposure. Using multi-omics and *in silico* modeling, we found that dormant bacteria reprogram metabolic pathways to modulate persister awakening. Accumulation of L-serine and reduction of arginine drive rapid resuscitation. L-serine promotes cysteine biosynthesis and motility while reducing energy metabolism to facilitate rapid resuscitation. In contrast, arginine slows regrowth from dormancy by enhancing ethanol-aldehyde and energy metabolism. L-serine and arginine can, respectively, promote or inhibit the regrowth of antibiotic persister cells in macrophages and mouse models, and regulate the awakening speed of *Salmonella*, *E. coli*, and methicillin-resistant *Staphylococcus aureus* (MRSA). These findings suggest new strategies to target chronic bacterial infections.

**Teaser:** L-serine speeds and arginine slows the awakening of antibiotic persisters, revealing targets for chronic infection.

## Introduction

Living cells regulate metabolic pathways to obtain energy from nutrients and control growth rate in response to environmental conditions. When cells encounter unfavorable environments, they arrest their overall metabolism(*1*). This phenomenon is widespread, observed across various organisms from bacteria to humans(*2*). When bacteria confront harsh environments, they produce dormant subpopulations as a survival strategy by arresting their metabolism(*3*). Dormant bacteria halt their metabolism to endure stresses such as antibiotics, host immune responses, and nutritional starvation(*4–6*). Notably, they can resuscitate their growth once favorable conditions return(*4–6*). Here, we investigate that dormant bacteria selectively control specific metabolic pathways, rather than overall metabolism, to regulate the rate of regrowth from dormancy. This promotes a paradigm shift regarding dormant cells, as dormant bacteria are currently understood to halt their overall metabolism.

Central carbon metabolism uses a complex set of enzymes to convert carbon sources into metabolic precursors and cellular enerygy(*7*). ATP and GTP are produced through central carbon metabolism such as glycolysis, the TCA cycle, and oxidative phosphorylation(*8*). In addition, precursors for cellular membranes and nucleic acids are produced by carbon metabolism (*9*). Thus, central carbon metabolism is the key regulatory pathway for cellular growth in response to environmental conditions(*7*). When cells encounter carbon starvation, amino acid metabolism can link with central carbon metabolism to continuously promote cellular growth(*10*). For instance, arginine and glutamic acid contribute to central carbon metabolism by supplying intermediates via glycolytic shunt pathways(*11, 12*). Additionally, serine enters the glycolysis pathway by converting to pyruvate(*13, 14*). During infection, bacteria can encounter carbon-limited conditions(*15, 16*), and thus may alter their central carbon metabolism to survive and regulate pathogenesis in the host. However, it is poorly understood how antibiotic persister bacteria regulate their central carbon metabolism during infections.

Dormant bacteria are highly tolerant of antibiotics and are called antibiotic persisters(*17*). Cellular metabolism of persister bacteria is arrested, resulting in evasion from antibiotic killing(*18*). They can resuscitate after antibiotic removal, thus causing chronic infections(*19*). The intracellular bacterium *Salmonella enterica* encounters unfavorable conditions within macrophage phagosomes, then forms persister subpopulations that evade antibiotic treatment and the immune system(*17*). Furthermore, bacteria can simultaneously produce a persister subpopulation under normal growth conditions, without stress, to prepare for harsh conditions such as antibiotic treatment (*20*). Understanding the regulatory mechanisms governing persister bacteria’s metabolism is critical for developing strategies to overcome chronic bacterial infections. This study elucidates the regulatory mechanism of metabolic pathways in antibiotic persisters, highlighting a novel strategy by which pathogens selectively control their metabolism to control resuscitation speed from persistence during infections.

In this study, we report that antibiotic persister bacteria selectively reprogram metabolism to control regrowth speed from dormancy during infection. To our knowledge, we systematically investigate, for the first time, the genome-wide functional metabolic capabilities of dormant bacterial populations with different regrowth speeds from dormancy through an integrative analysis based on metabolomics, proteomics, and *in silico* modeling. Dormant bacterial subpopulations accumulate L-serine and reduce arginine, promoting rapid resuscitation. For rapid resuscitation from dormancy, L-serine activates cysteine biosynthesis and motility, while inhibiting energy metabolism. By contrast, arginine reduces translation and motility, induces ethanol-aldehyde and energy metabolism, and thereby extends dormancy. Notably, L-serine and arginine regulate the resuscitation of antibiotic persister bacteria, a type of dormant bacteria under antibiotic treatment, in macrophages and mouse models. In addition, L-serine and arginine control the resuscitation of various antibiotic-persistent bacteria, including *Salmonella enterica*, *Escherichia coli*, and a clinically isolated methicillin-resistant *Staphylococcus aureus*. Our findings demonstrate pathogens’ ability to selectively regulate metabolic pathways responding to unfavorable conditions, including antibiotic treatment, to speed return to the regrowth stage, illuminating critical aspects of host reinfection and chronic infections.

## Results

### Dormant bacteria differentially regulate metabolic pathways to control regrowth speed

When bacteria encounter unfavorable conditions, such as antibiotics and nutrient starvation, they enter dormant states(*21*). Dormant bacteria reduce ATP production(*22*) and attempt to reduce protein degradation(*23*). This mechanism is critical for rapid regrowth from dormancy(*24*). To date, dormant bacteria are known to arrest their overall metabolism to reduce ATP levels, which enables bacteria to enter dormancy(*21*). However, it remains poorly understood whether dormant bacteria arrest their entire metabolism or selectively regulate metabolic pathways during dormancy. Furthermore, it is not clear which metabolic pathways during dormancy control resuscitation in bacteria.

To address this question, we conducted proteomic and metabolomic analyses to identify the metabolic pathways regulated during dormancy to promote rapid regrowth in the pathogenic bacterium *Salmonella enterica*. We previously reported that dormant bacteria treated with sublethal bacteriostatic antibiotic chloramphenicol (10 μg/ml) exhibit slow regrowth from dormancy, attributed to impaired ATP reduction resulting from metabolic arrest during dormancy(*6*). Fast-regrowing dormant bacteria exemplify a natural dormant state, characterized by reduced ATP levels during dormancy due to metabolic arrest(*6*). Therefore, we tested bacteria in growth-inhibiting conditions that either increased or decreased ATP levels, and subsequently assessed bacterial regrowth upon transfer to fresh media(*6*). To investigate the proteome and metabolome profiles of fast– and slow-regrowth dormant bacteria, we collected samples at multiple time points: starting growth (a and b), dormant states (c and d), preparing for regrowth from dormancy in fresh medium (e and f), and actively regrowing state (g and h) (Fig. 1A and B). These samples were subjected to liquid chromatography-mass spectrometry for proteomic and metabolomic profiling.

**Figure 1.**
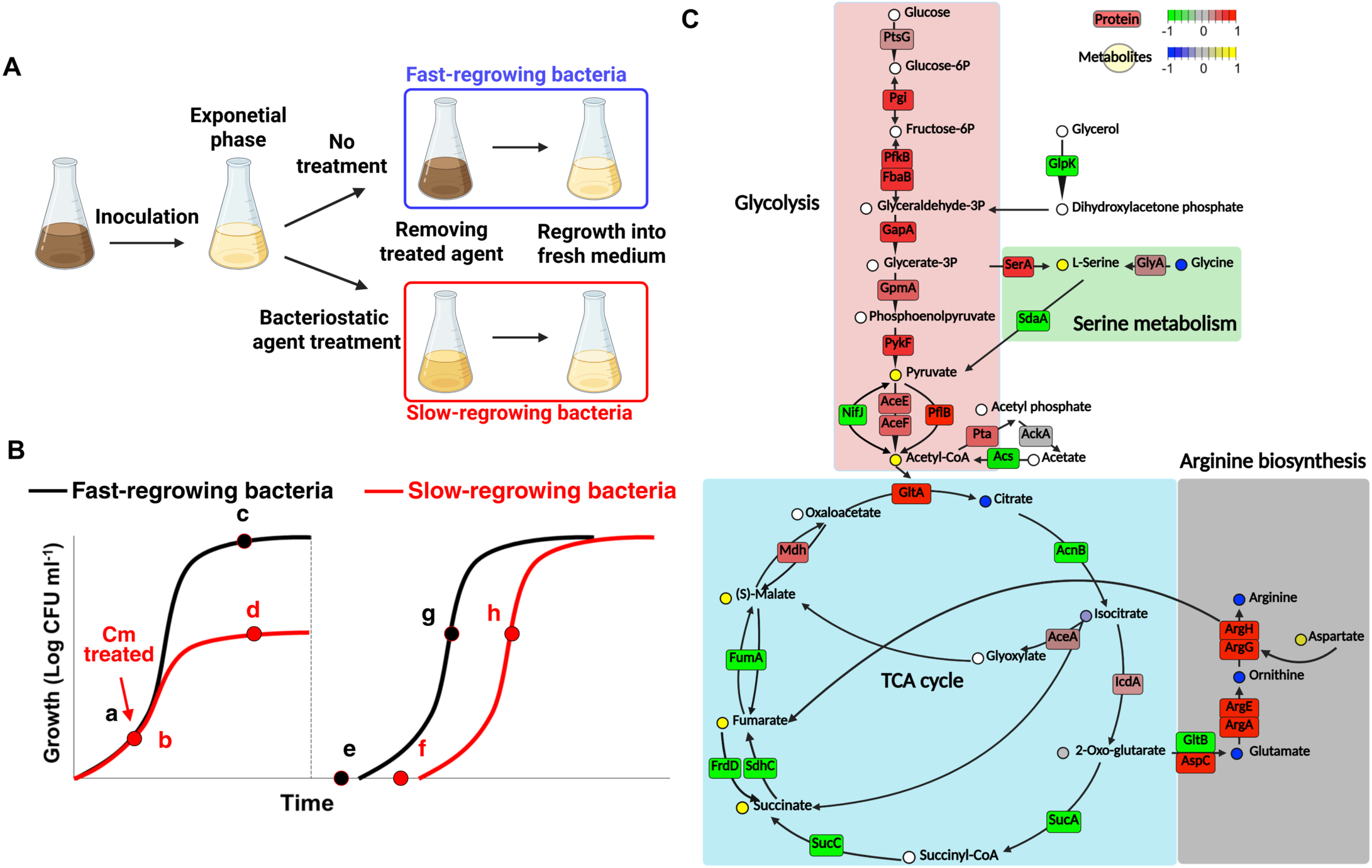
Dormant bacteria selectively regulate their metabolic pathways to control regrowth from dormancy. (A) Schematic of the regrowth assay in fast– or slow regrowing bacteria from dormancy. Bacteria were grown in N-minimal media pH 7.7 containing 1mM MgCl_2_ for 16 h. Then, bacteria were diluted into fresh N-minimal media and grown to mid-exponential phase for 2.5 h. And then, bacteria cultures were split and either exposed to chloramphenicol (Cm, 20 μg/mL) to induce slow-regrowing dormancy or left untreated to induce fast-regrowing dormancy(*6*). Each culture was then incubated to its stationary-phase plateau for 3 h. After washing to remove Cm, same number bacteria by normalization of OD_600_ values were reinoculated into fresh N-minimal media to monitor regrowth. Untreated group showed rapid regrowth (fast-regrowing bacteria, blue box), whereas Cm-pretreated group exhibited delayed recovery (slow-regrowing bacteria, red box). (B) Schematic of growth curves of fast-regrowing (black line) and slow-regrowing (red line) dormant bacteria during the initial growth phase (a–d) for dormancy and following reinoculation (e–h) for regrowth. (C) Central metabolic pathways reconstructed from integrative proteomic and metabolomic analyses comparing fast-regrowing bacteria (c in panel B) and slow-regrowing bacteria (d in panel B) during dormancy. Circles represent metabolites and squares represent proteins. Color indicates relative abundance in fast-regrowing dormant bacteria (c in panel B) relative to slow-regrowing dormant bacteria (d in panel B): red (increased protein amounts in fast-regrowing dormant bacteria), green (decreased protein amounts in fast-regrowing dormant bacteria), yellow (increased metabolite abundance in fast-regrowing dormant bacteria), and blue (decreased metabolite abundance in fast-regrowing dormant bacteria). Glycolysis (pink box) and serine metabolism (green box) are upregulated in fast-regrowing dormant bacteria during dormancy, whereas the TCA cycle (blue box) and arginine biosynthesis (grey box) are downregulated in fast-regrowing dormant bacteria during dormancy.

Multiple lines of evidence suggest that bacteria differentially regulate metabolic pathways during dormancy, rather than overall metabolism for rapid regrowth. First, proteins related to energy production pathways, such as the TCA cycle and oxidative phosphorylation, significantly decreased in fast-growing dormant bacteria (Fig. S1A). Second, protein synthesis pathways, including RNA stabilization, translation, rRNA synthesis, and ribosome synthesis, also decreased substantially in fast-growing dormant bacteria (Fig. S1A). Third, proteins related to amino acid synthesis, nutrient transportation, stress response, and cell wall synthesis pathways increased in fast-growing dormant cells (Fig. S1A). Fourth, metabolites related to TCA cycles, phospholipid synthesis, purine metabolism, pyrimidine metabolism, and vitamin metabolism significantly accumulated in fast-regrowing dormant populations (Fig. S1B), because amounts of metabolic enzymes related to above significantly were decreased (Fig. S1A). Lastly, amino acids except for serine decreased in fast-regrowing dormant bacteria compared to slow-regrowing bacteria (Fig. S1B), because amounts of metabolic enzymes related to above were increased (Fig. S1A). Together, dormant bacteria selectively regulate their metabolism thereby regrowing rapidly from dormancy.

Furthermore, we conducted *in silico* modeling for global metabolite flux simulations in *E. coli* during dormancy to predict the change of metabolic pathways in fast– or slow-regrowing bacteria from dormancy. For predictive modeling for metabolic flux analysis, slow-regrowing dormant bacteria exhibit high ATP levels, whereas fast-regrowing dormant bacteria have low ATP levels during dormancy. Consistent with results from metablome and proteome analysis, *in silico* modeling reveals significant activation of glycolysis, acetate synthesis, serine metabolism, and aspartate-fumarate metabolism in fast-regrowing dormant bacteria characterized by low ATP levels (Fig. S2A). By contrast, the metabolic flux of the TCA cycle and the pyruvate-malate metabolic pathway is significantly elevated in slow-regrowing dormant bacteria exhibiting high ATP levels (Fig. S2B). In conclusion, integrative profiling analysis and *in silico* metabolic flux modeling indicate that bacteria selectively regulate their metabolism during dormancy, which is essential for determining the regrowth speed of dormant bacteria.

### Central carbon metabolism is reprogrammed between fast– and slow-regrowing dormant bacteria

To examine which metabolites are essential for the rapid resuscitation of dormant bacteria, we integrated proteome and metabolome data for fast– or slow-regrowing dormant bacteria (Fig. 1C). Interestingly, central carbon metabolism, which is a vital metabolic pathway for cellular growth and survival(*7*), is significantly different between fast– and slow-regrowing dormant bacteria (Fig. 1C). We employ a color code in metabolic pathways (Fig. 1C) to illustrate the disparity in central carbon metabolism between fast-regrowing dormant bacteria (denoted as c in Fig. 1B) and slow-regrowing dormant bacteria (denoted as d in Fig. 1B). Enhanced and reduced proteins in fast-regrowing dormant bacteria were indicated by red and green colors, respectively, compared to slow-regrowing dormant bacteria. In addition, metabolites accumulated and diminished in fast-regrowing dormant bacteria formed a yellow circle, whereas those in slow-regrowing dormant bacteria formed a blue circle.

It is well known that central carbon metabolism, which comprises glycolysis and the TCA cycle, utilizes carbon sources to produce energy and structural components for cellular constituents(*9*). Additionally, the synthetic and catabolic pathways of amino acids can influence central carbon metabolism(*12, 25*). Thus, we hypothesized that dormant bacteria regulate the central carbon metabolism and related amino acid metabolism to promote rapid resuscitation from dormancy. Multiple lines of evidence support the notion that fast– and slow-regrowing dormant bacteria differentially regulate central carbon metabolism. First, enzymes related to the glycolysis pathway were highly accumulated in fast-regrowing dormant bacteria (Fig. 1C). Second, enzymes for the TCA cycle were dramatically reduced (Fig. 1C). Third, several enzymes involved in serine and arginine metabolism, which are connected with central carbon metabolism, were highly increased or decreased in rapidly regrowing dormant bacteria (Fig. 1C). Fourth, several metabolites, including serine, pyruvate, fumarate, and succinate, significantly accumulated in fast-regrowing dormant cells (Fig. 1C). By contrast, arginine, glutamate, aspartate, citrate, and ornithine decreased in fast-resuscitating dormant bacteria (Fig. 1C). Together, dormant bacteria selectively regulate enzymes and metabolites in central carbon metabolism, which may control bacterial resuscitation from dormancy.

### The accumulation of L-serine promotes rapid resuscitation of dormant bacteria

During infection, bacterial pathogens frequently encounter carbon starvation(*15, 16*). Bacteria can enter a dormant state under carbon starvation conditions, and amino acid metabolism can link with central carbon metabolism to continuously promote adaptation and proliferation(*10*). For instance, amino acids act as signals for latent spores in Gram-positive bacteria(*26*). In addition, our data indicate that several amino acids involved in central carbon metabolism are altered in rapidly regrowing dormant bacteria (Fig. 1). Thus, we hypothesized that these metabolites could serve as signals or essential nutrients for rapid bacterial resuscitation from dormancy.

During dormancy, *Salmonella* exhibits a significant increase in L-serine amounts due to augmented levels of the synthesizing enzyme SerA and diminished levels of the degrading enzyme SdaA in fast-regrowing dormant bacteria (Fig. 2A). Consistently, fast-regrowing dormant bacteria exhibited elevated amounts of SerA protein (Fig. 2B) and diminished amounts of SdaA proteins (Fig. 2C) compared to slow-regrowing dormant bacteria. Furthermore, *in silico* modeling indicated that the serine synthetic pathway is significantly activated by the glycerate-3P metabolite of glycolysis in fast-regrowing dormant bacteria (Fig. 2D), but not in slow-regrowing dormant bacteria (Fig. 2E).

**Figure 2.**
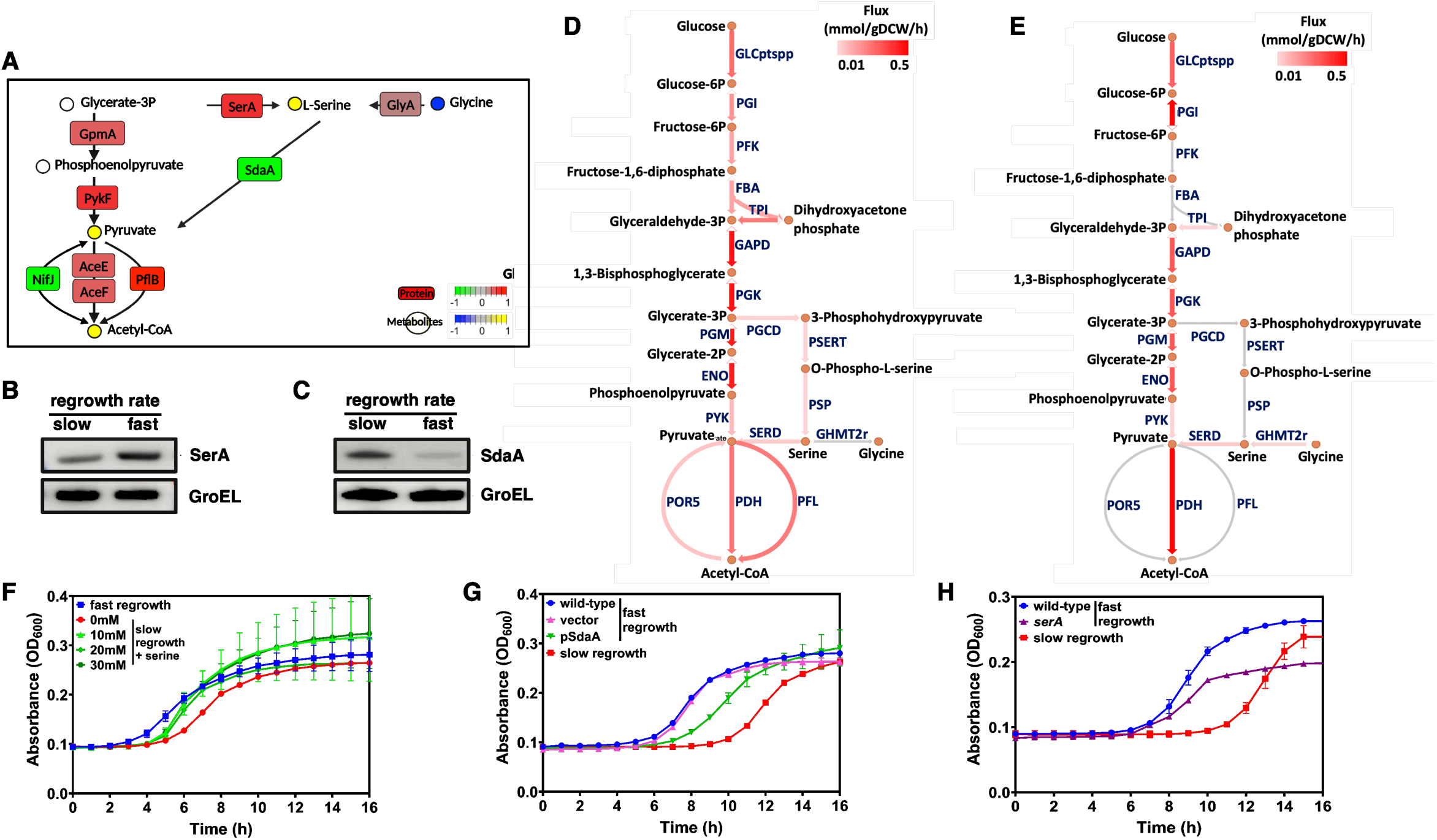
L-serine facilitates rapid resuscitation from dormancy in *Salmonella*. (A) Metabolic map of serine biosynthesis and degradation based on integrative proteomics and metabolomics analyses comparing fast-regrowing and slow-regrowing dormant bacteria. Circles represent metabolites and squares represent proteins. Color indicates relative abundance in fast-regrowing dormant bacteria relative to slow-regrowing dormant bacteria: red (increased protein amounts in fast-regrowing dormant bacteria), green (decreased protein amounts in fast-regrowing dormant bacteria), yellow (increased metabolite abundance in fast-regrowing dormant bacteria), and blue (decreased metabolite abundance in fast-regrowing dormant bacteria). (B and C) Immunoblot analysis determines the amounts of SerA (B) and SdaA (C) proteins during dormancy in *S*. Typhimurium ATCC 14028s. Bacteria were grown in N-minimal media pH 7.7 containing 1mM MgCl2 for 16 h. After that, bacteria were diluted into fresh N-minimal media and grown to mid-exponential phase for 2.5 h. And then, bacteria were grown for 3 h after treatment with chloramphenicol (20 μg/mL) to produce slow-regrowing dormant bacteria or without chloramphenicol to induce fast-regrowing dormant bacteria. (B and C) Immunoblot analysis showing that SerA, a serine biosynthesis enzyme, is enriched (B) and SdaA, a serine degradation enzyme, is reduced (C) in fast-regrowing dormant bacteria. GroEL served as a loading control. (D-E) *In silico* metabolic flux analysis reveals differential regulation of serine metabolic pathways between fast– and slow-regrowing dormant bacteria. Metabolic flux comparison between fast-regrowing dormant bacteria (low-ATP) (D) and slow-regrowing dormant bacteria (high-ATP) (E) groups within serine biosynthesis and degradation pathways, because high ATP promotes rapide resuscitation of dormant bacteria(*6*). Reactions are depicted as arrows with abbreviation labels in blue, and metabolites are shown as orange circles. Reactions carrying no flux are shown in grey, and both the color intensity and arrow width indicate the flux magnitude in mmol/gDCW/h. Abbreviation for a metabolic enzyme. ENO, Enolase; FBA, Fructose-bisphosphate aldolase; GAPD, Glyceraldehyde-3-phosphate dehydrogenase; GHMT2r, Glycine hydroxymethyltransferase reversible; GLCptspp, D-glucose transport via PEP:Pyr PTS; PDH, Pyruvate dehydrogenase; PFK, Phosphofructokinase; PFL, Pyruvate formate lyase; PGCD, Phosphoglycerate dehydrogenase; PGI, Glucose-6-phosphate isomerase; PGK, Phosphoglycerate kinase; PGM, Phosphoglycerate mutase; POR5, Pyruvate synthase; PSERT, Phosphoserine transaminase; PSP, Phosphoserine phosphatase; PYK, Pyruvate kinase; SERD, L-serine deaminase; TPI, Triose-phosphate isomerase. (F-H) Regrowth analysis of dormant bacteria in serine treatment (F), SdaA overexpression (G), and *serA* mutant strain (H). Overnight cultures of *S*. Typhimurium ATCC 14028s were grown into fresh N-minimal medium for 2.5 h and then treated with chloramphenicol (20 μg/mL) for 3 h to induce slow-regrowing dormant bacteria. Also, overnight cultures were grown into fresh N-minimal medium at 37 °C for 5.5 h to produce fast-regrowing dormant bacteria. After washing to remove the antibiotic, bacteria were re-inoculated in fresh N-minimal medium, then bacterial regrowth was monitored at 37 °C for 16 h using a microplate reader. (F) Bacterial regrowth was monitored at 37 °C for 16 h using a microplate reader in the presence of 0, 10, 20, or 30 mM L-serine. Supplementation with L-serine accelerated the regrowth of dormant bacteria in a dose-dependent manner. (G) Bacterial regrowth was monitored at 37 °C for 16 h with wild-type harboring plasmid with/without *sdaA* gene. Expression of *sdaA*from a heterologous promoter in a plasmid delays bacterial regrowth from dormancy. (H) Bacterial regrowth was monitored at 37 °C for 16 h with wild-type and *serA* mutant strains. Inactivation of *serA* results in delayed regrowth under fast regrowth conditions.

Consistent with the discovery that fast-regrowing dormant bacteria accumulate L-serine, we reasoned that L-serine may facilitate the rapid resuscitation of dormant bacteria. L-serine treatment markedly enhanced the regrowth rate relative to slow-regrowing dormant bacteria (Fig. 2F). By contrast, D-serine treatment did not influence regrowth rate in dormant bacteria (Fig. S3A). The expression of the L-serine-degrading enzyme SdaA under a heterologous promoter led to a gradual resuscitation of fast-regrowing dormant bacteria (Fig. 2G) through serine reduction (Fig. S3B). In contrast, the resuscitation rate with the empty plasmid was comparable to that of the wild-type (Fig. 2G). In addition, the inactivation of the *serA* gene, responsible for encoding the L-serine-producing enzyme, diminished the regrowth of fast-regrowing dormant bacteria (Fig. 2H) because of serine depletion (Fig. S3C). Furthermore, to investigate whether L-serine acts as a signal to induce rapid resuscitation from dormancy, we treated L-serine analog alpha-methyl-L-serine, which is not metabolized by bacteria (*27*), to dormant bacteria. L-serine acts as a signal to induce rapid resuscitation, because alpha-methyl-L-serine treatment increased the rapid resuscitation of dormant bacteria (Fig. S3D). Collectively, these findings indicate that the accumulation of L-serine facilitates the resuscitation of dormant bacteria.

### L-serine facilitates rapid regrowth from dormancy by decreasing energy production, improving stress defense mechanisms and stimulating motility

To investigate how L-serine regulates the resuscitation of dormant bacteria, we conducted proteome profiling of slow-regrowing dormant bacteria with and without L-serine treatment throughout the lag and log phases of regrowth from dormancy (Fig. 3A).

**Figure 3.**
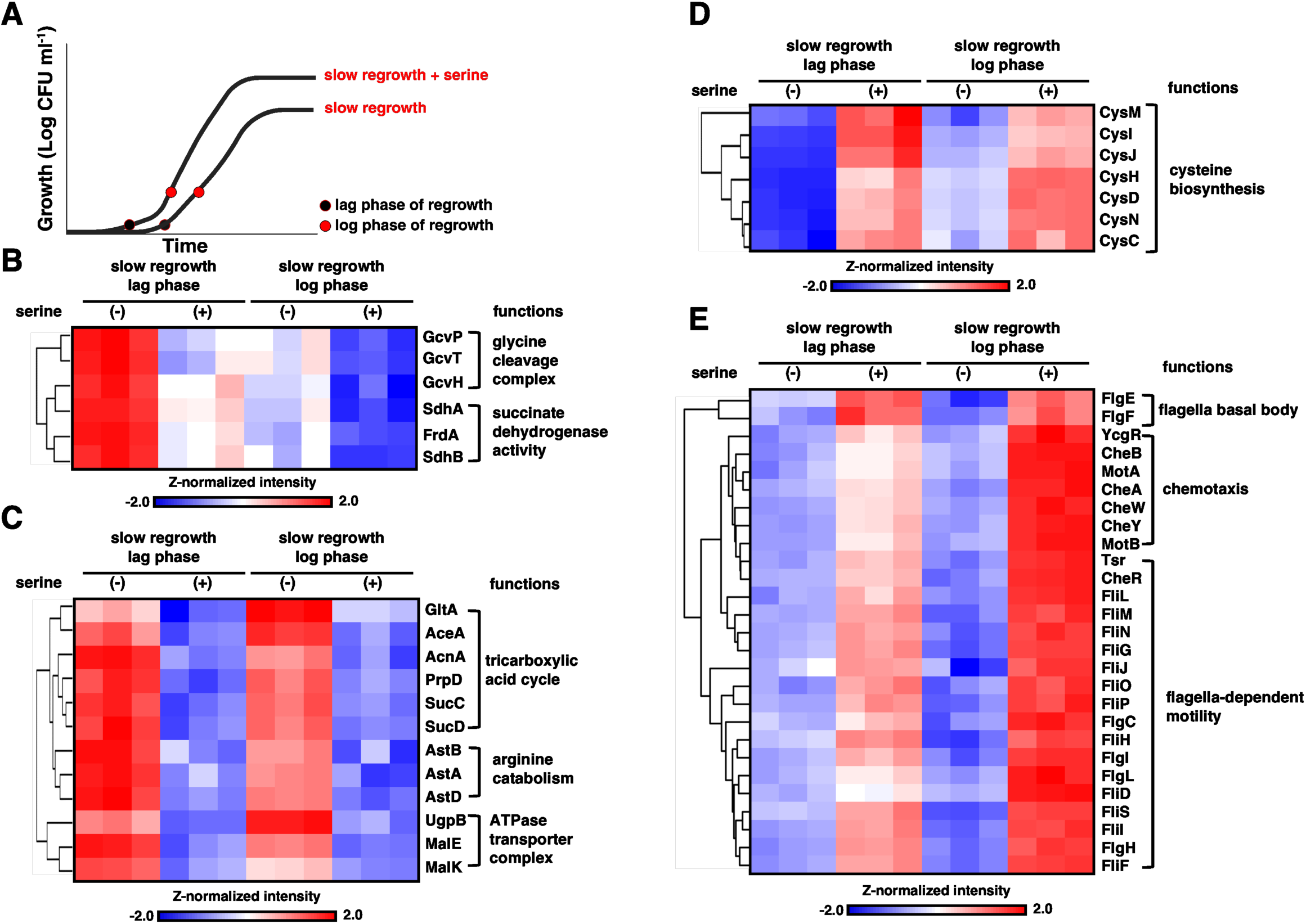
L-Serine facilitates dormancy exit by selectively reprogramming the proteome. (A) Schematic of the regrowth from dormancy with the supplement of L-serine. Bacteria were grown in N-minimal media, pH 7.7, containing 1mM MgCl_2_ for 16 h. Then, bacteria were diluted into fresh N-minimal media and grown to mid-exponential phase for 2.5 h. And then, bacterial cultures were exposed to chloramphenicol (Cm, 20 μg/mL) to produce slow-regrowing dormant bacteria. Culture was then incubated in its stationary-phase plateau for 3 h. After washing to remove Cm, the same number of bacteria, normalized by OD_600_, were reinoculated into fresh N-minimal media to monitor regrowth in the presence or absence of 30 mM L-serine. (B–E) Heatmaps via hierarchical cluster analysis (HCA) show proteomic changes in regrowing dormant *S*. Typhimurium bacteria from each growth point in panel A. (B and C) Proteins that were significantly enriched in the slow-regrowing dormant bacteria compared to slow-regrowing dormant bacteria with L-serine at the lag phase of regrowth. These proteins were either down-regulated during the log phase (B) or remained elevated in the lag and log phases (C), indicating selective retention of a dormancy-associated proteome in slowly regrowing dormant bacteria. (D and E) Proteins that were significantly upregulated in slow-regrowing dormant bacteria with L-serine compared to slow-regrowing dormant bacteria at the lag phase of the regrowth. These early serine-responsive proteins were either continuously induced in lag or log phases (D). Several proteins up-regulated by L-serine treatment continued to increase in amount through log phase (E).

The glycine cleavage complex and succinate dehydrogenase pathways are significantly activated when slow-regrowing dormant bacteria transition to active regrowth (Fig. 3B). In contrast, these pathways remain inactive during the lag phase of regrowth in slow-regrowing dormant bacteria (Fig. 3B). Notably, L-serine treatment of slow-regrowing dormant bacteria suppressed expression of both the glycine cleavage complex and succinate dehydrogenase pathways during regrowth (Fig. 3B). Glycine cleavage is essential for energy production through NADH and acetyl-CoA generation(*28, 29*), and succinate dehydrogenase enzymes contribute to energy production via the TCA cycle(*30*). Given that fast-regrowing dormant bacteria exhibit reduced energy production (Fig. S1A), L-serine accumulation appears to attenuate activation of the energy production pathway, potentially preventing energy wastage during the initial stages of dormancy exit. Furthermore, while slow-regrowing dormant bacteria displayed increased levels of the TCA cycle, arginine catabolism, and the glycerol-3-phosphate (G3P) transporter system during regrowth, treatment with L-serine resulted in a reduction of these proteins (Fig. 3C). Because the TCA cycle, arginine catabolism, and G3P transporters are crucial to cellular energy production(*31, 32*), their reduction by L-serine treatment in slow-regrowing dormant bacteria promotes rapid regrowth from dormancy by reducing unnecessary energy expenditure. This corresponds to our previous findings that fast-regrowing dormant bacteria reduced energy production pathways during dormancy (Fig. 1 and Fig. S1A). Thus, L-serine reduces protein levels associated with the TCA cycle, arginine catabolism, and the glycerol-3-phosphate (G3P) transporter system during regrowth, thereby facilitating rapid regrowth.

Next, we investigated which pathways are increased by L-serine treatment during regrowth from dormancy. The synthesis of cysteine is crucial for addressing oxidative stress and antibiotic resistance(*33*). Also, the accumulation of oxidative stress-related proteins in fast-regrowing dormant bacteria (Fig. S1A). In agreement with this, the activation of the cysteine biosynthesis pathway through L-serine treatment facilitates swift regrowth from dormancy (Fig. 3D). Additionally, bacteria conserve energy by reducing motility and flagella production under nutrient-limited conditions(*34*). Flagellar-dependent motility and chemotaxis facilitate bacterial adaptation to new environments by improving access to resources and aiding in the avoidance of unfavorable conditions(*34*). Thus, L-serine treatment significantly enhances flagellar-dependent movement and chemotaxis, potentially indicating rapid resuscitation of dormant bacteria (Fig. 3E). Collectively, L-serine promotes the rapid regrowth of slowly regrowing dormant bacteria by diminishing energy production, activating oxidative stress defense mechanisms, and enhancing bacterial movement, such as motility and chemotaxis.

### Reduction of arginine and production of fumarate facilitate resuscitation of dormant bacteria

Subsequently, we investigated which metabolites delay the regrowth of dormant bacteria. Interestingly, proteome profiling study indicated a significant increase in arginine synthesis enzymes in fast-regrowing dormant bacteria (Fig. 4A). The amounts of ArgH protein are higher in fast-regrowing dormant bacteria compared to slow-regrowing dormant bacteria (Fig. 4B). The synthesis of enzymes increased in fast-regrowing dormant bacteria, while the levels of arginine significantly decreased in these cells, as indicated by metabolome profiling (Fig. 4A). The arginine biosynthesis pathway is also implicated in fumarate production (Fig. 4A). Furthermore, *in silico* modeling demonstrated that aspartate is converted to fumarate, rather than arginine, in fast-regrowing dormant bacteria (Fig. 4C). In contrast, conversion pathway from the aspartate to fumarate is not activated in slow-regrowing dormant bacteria. (Fig. 4D). Thus, we hypothesized that the reduction of arginine and the elevation of fumarate may facilitate the rapid resuscitation of dormant bacteria.

**Figure 4.**
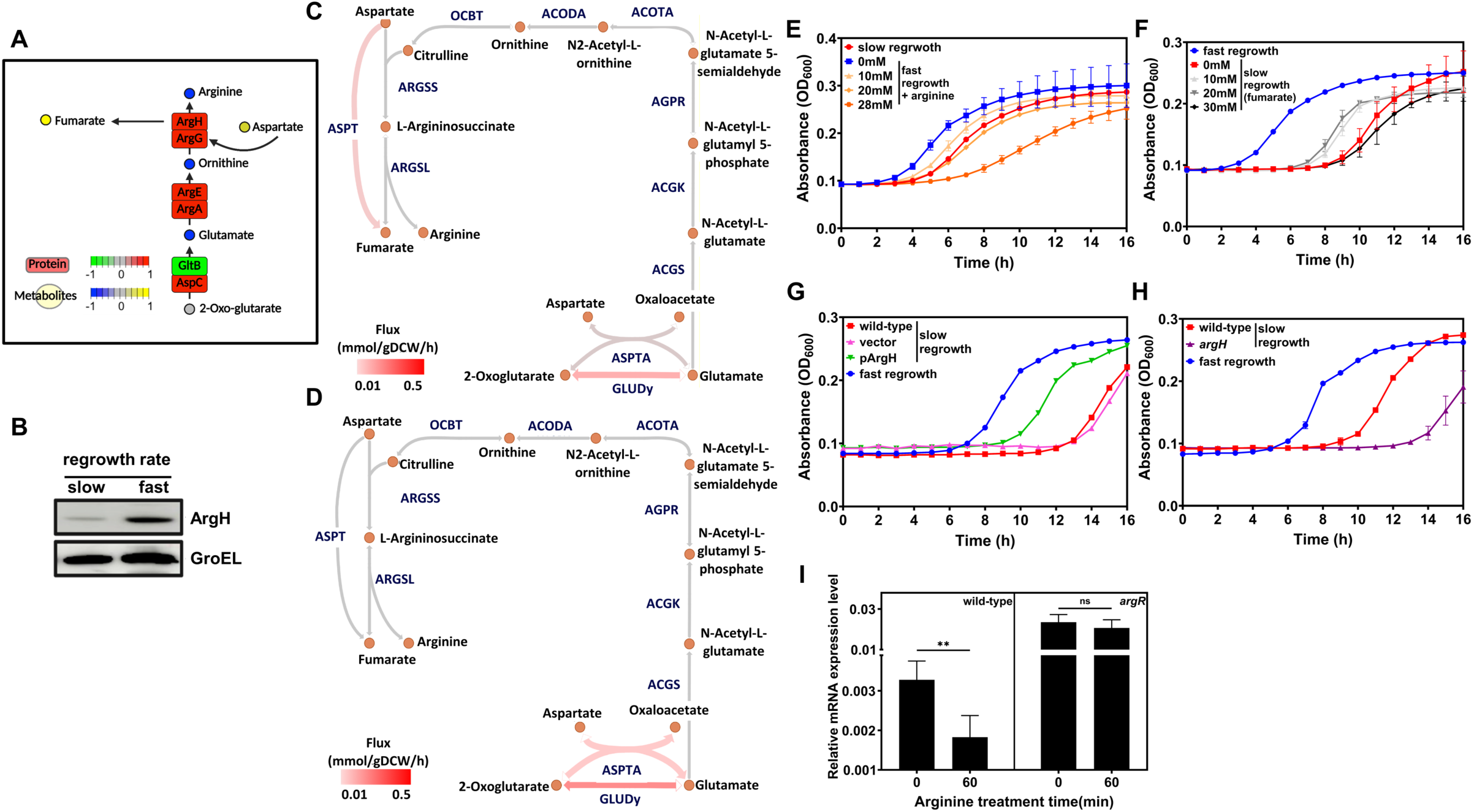
Arginine delays bacterial regrowth from dormancy by inhibiting fumarate synthesis. (A) Metabolic map of arginine biosynthesis pathway based on proteomics and metabolomics analyses comparing fast– and slow-regrowing dormant bacteria. Circles represent metabolites and squares represent proteins. Color indicates relative abundance in fast-regrowing bacteria relative to slow-regrowing dormant bacteria: red (increased protein amounts in fast-regrowing dormant bacteria), green (decreased protein amounts in fast-regrowing dormant bacteria), yellow (increased metabolite abundance in fast-regrowing dormant bacteria), and blue (decreased metabolite abundance in fast-regrowing dormant bacteria). (B) Immunoblot analysis showing that ArgH, which catalyzes the conversion of argininosuccinate to arginine and fumarate, is enriched in fast-regrowing dormant bacteria. Bacteria were treated with or without chloramphenicol (20 μg/mL) for 3 h to induce dormancy. GroEL served as a loading control. (C and D) *In silico* metabolic flux analysis was performed to compare fast– and slow-regrowing dormant bacteria in arginine metabolic pathways. Metabolic flux comparison between fast-regrowing dormant bacteria (low-ATP) (C) and slow-regrowing dormant bacteria (high-ATP) (D) groups within arginine biosynthesis pathways, because high ATP promotes rapid resuscitation of dormant bacteria(*6*). Reactions are depicted as arrows with abbreviation labels in blue, and metabolites are shown as orange circles. Reactions carrying no flux are shown in grey, and both the color intensity and arrow width indicate the flux magnitude in mmol/gDCW/h. Abbreviation for a metabolic enzyme. ACGK, Acetylglutamate kinase; ACGS, N-acetylglutamate synthase; ACODA, Acetylornithine deacetylase; ACOTA, Acetylornithine transaminase; AGPR, N-acetyl-g-glutamyl-phosphate reductase; ARGSL, Argininosuccinate lyase; ARGSS, Argininosuccinate synthase; ASPT, L-aspartase; ASPTA, Aspartate transaminase; GLUDy, Glutamate dehydrogenase; OCBT, Ornithine carbamoyltransferase. (E-H) Regrowth analysis of dormant bacteria in arginine treatment (E), arginine treatment (F), ArgH overexpression (G), and *argH* mutant strain (H). Overnight cultures of *S*. Typhimurium ATCC 14028s were grown into fresh N-minimal medium for 2.5 h and treated with chloramphenicol (20 μg/mL) for 3 h to induce slow-growing dormant bacteria. Also, overnight cultures were grown into fresh N-minimal medium at 37 °C for 5.5 h to produce fast-regrowing dormant bacteria. After washing to remove the antibiotic, bacteria were re-inoculated in fresh N-minimal medium, then bacterial regrowth was monitored at 37 °C for 16 h using a microplate reader.(E) Bacterial regrowth was monitored at 37 °C for 16 h using a microplate reader in the presence of 0, 10, 20, or 30 mM L-arginine. Exogenous arginine delayed regrowth in a concentration-dependent manner. (F) Bacterial regrowth was monitored at 37 °C for 16 h using a microplate reader in the presence of 0, 10, 20, 30 mM fumarate. Supplementation with fumarate promotes rapid regrowth in slow-regrowing dormant bacteria. (G) Bacterial regrowth was monitored at 37 °C for 16 h with wild-type harboring plasmid with/without *argH* gene. Expression of *argH* from a heterologous promoter in a plasmid accelerates bacterial regrowth from dormancy. (H) Bacterial regrowth was monitored at 37 °C for 16 h with wild-type and *argH* mutant strains. Inactivation of *argH* results in delayed regrowth under slow regrowth conditions. (I) The mRNA amount of argH under treatment with arginine indicates that *argH* transcription is downregulated following arginine treatment in wild-type bacteria but not in the *argR* mutant. Unpaired Student’s t tests were performed between 0 and 60 min arginine treatment within each strain; \*\**P* < 0.01, ns, not significant. The mean and SD from three independent experiments are shown.

Multiple lines of evidence support the notion that the reduction of arginine and the accumulation of fumarate promote the resuscitation of dormant bacteria. First, treatment with arginine resulted in a decreased regrowth rate of fast-regrowing dormant bacteria (Fig. 4E). Second, treatment of exogenous fumarate to slow-regrowing dormant bacteria increased the regrowth speed from dormancy (Fig. 4F). By contrast, other metabolites in the arginine biosynthetic pathway, such as glutamate and aspartic acid, did not change the regrowth speed of dormant bacteria (Fig. S4). Third, overexpression of *argH* by heterologous promoters in plasmids led to faster regrowth of slow-regrowing dormant bacteria (Fig 4G) because the amounts of fumarate were increased (Fig. S5A). In contrast, the overexpression of *argH* via heterologous promoters in plasmids does not alter arginine levels (Fig. S5B). Fourth, inactivation of *argH* resulted in slower resuscitation of slow-regrowing bacteria (Fig. 4H) due to fumarate reduction (Fig. S5C). Also, *argH* inactivation did not change arginine levels in slow-regrowing bacteria (Fig. S5D). Lastly, because ArgR represses *argH* expression via arginine binding(*35*), arginine inhibits production of fumarate by decreasing ArgH amounts that confers slow regrowth of dormant bacteria (Fig. 4I). Together, reduction of arginine and production of fumarate are required for fast resuscitation of dormant bacteria.

### Arginine inhibits dormant exit by suppressing ribosome rebuilding and flagella synthesis, while enhancing energy production and ethanol-aldehyde metabolism

To investigate the mechanism of arginine on bacterial regrowth delay from dormancy, we conducted a proteome profiling analysis of fast-regrowing dormant bacteria in the presence of arginine (Fig. 5A). First, arginine treatment decreased the synthesis and assembly of ribosomes during the transition of fast-regrowing dormant bacteria to active regrowth (Fig. 5B). According to previous proteome profiling results (Fig. S1A), the levels of ribosomal components are anticipated to remain low during dormancy to enable swift regrowth. Rapid ribosome synthesis is likely required for rapid regrowth from dormancy (Fig. 5B), facilitating the production of various proteins that promote accelerated biological activity. In agreement with this, arginine prolonged the dormant state by reducing ribosome reconstruction (Fig. 5B). Second, arginine significantly reduced flagella biosynthesis, motility, and chemotaxis (Fig. 5C), contrasting sharply with the effects observed from L-serine treatment (Fig. 3E). Flagellar-dependent movement and chemotaxis indicate resuscitation from dormancy (Fig. 3E), as bacteria produce flagella to navigate favorable environments and avoid less favorable conditions following dormancy(*34*). Thus, arginine inhibits resuscitation from dormancy by decreasing flagella synthesis, motility, and chemotaxis (Fig. 5C).

**Figure 5.**
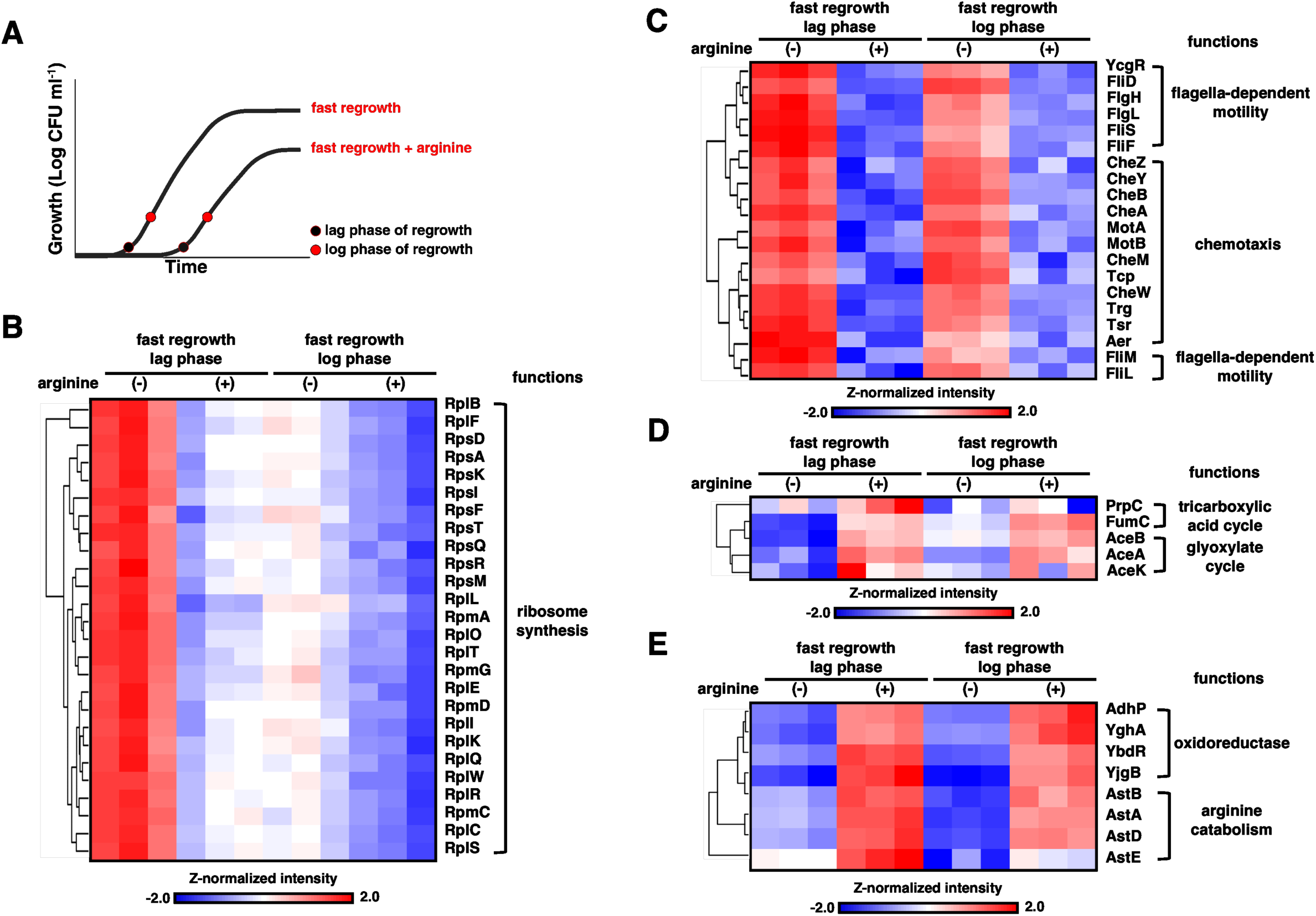
Arginine delays regrowth by selectively reprogramming the proteome. (A) Schematic overview of regrowth or dormant bacterin in the presence or absence of arginine. Fast-regrowing dormant bacteria were washed and transferred into fresh medium with or without L-arginine supplementation. Samples were harvested at lag and log regrowth phases. (B–E) Heatmaps via hierarchical cluster analysis (HCA) show proteomic changes in *S*. Typhimurium cells transferred into fresh media with or without L-arginine supplementation in fast-regrowing dormant bacteria. Samples were collected at two time points: the lag phase and the log phase. (B and C) Proteins that were significantly enriched in the arginine-untreated group (arginine–) compared to the arginine-treated group (arginine+) at the lag phase. These proteins were either downregulated (B) or remained elevated (C) during the log phase. (D and E) Proteins that were significantly upregulated in the arginine-treated group (arginine+) compared to the arginine– group at the lag phase. These proteins were either weakly induced (D) or strongly induced (E) during regrowth.

In contrast, arginine promoted energy production in fast-regrowing dormant bacteria during regrowth (Fig. 5D), which is precisely the opposite effect observed with L-serine treatment (Fig. 3C). Previous findings indicate that fast-regrowing dormant bacteria decrease energy production during dormancy and regrowth stage (Fig. 3C and Fig. S1A). The overactivation of energy production pathways, including the TCA cycle and glyoxylate cycle, via arginine can prolong the dormant condition (Fig. 5D). Moreover, arginine markedly increases the levels of aldehyde metabolism-related enzymes in fast-regrowing dormant bacteria, leading to an extended dormancy (Fig. 5E). The metabolism of ethanol and aldehyde may indicate the stationary phase of bacterial growth and low oxygen stress(*36*). According to this, arginine consistently induces enzymes involved in ethanol-aldehyde metabolism, thereby maintaining dormancy. Finally, as expected, the enzymes associated with arginine catabolism were increased after arginine treatment. Altogether, arginine treatment inhibits the regrowth of dormant bacteria by reducing ribosome synthesis, decreasing flagella-dependent motility, enhancing energy production systems, and activating ethanol-aldehyde metabolism.

### L-Serine and arginine govern the resuscitation speed of antibiotic-induced persisters

Bacterial dormancy promotes the formation of persister populations that enable bacteria to be tolerant of antibiotics without genetic alteration(*35*). Persister bacteria can resuscitate from dormancy following a reduction in antibiotic concentration, thereby contributing to chronic infections in the host(*37*). In this study, we determined that L-serine accumulation and arginine reduction facilitate the resuscitation of dormant bacteria (Fig. 2 and 4). Therefore, we reasoned that these amino acids might regulate the resuscitation of antibiotic persister bacteria

To test whether L-serine and arginine control regrowth rate of antibiotic persisters, we measured bacterial regrowth by absorbance (OD600) and colony forming units (CFUs) with exogenous L-serine or arginine added to antibiotic persisters, which were generated by treatment with the bactericidal antibiotics ciprofloxacin (Fig. 6A). Multiple lines of evidence support the notion that L-serine and arginine regulate resuscitation of antibiotic persisters during infections. First, L-serine treatment promotes the resuscitation of ciprofloxacin antibiotics-induced *Salmonella* persisters (Fig. 6B). By contrast, persister bacteria treated with arginine showed markedly slower resuscitation compared to untreated persister bacteria (Fig. 6B). Second, colony numbers for regrowing persisters rapidly increased with L-serine treatment (Fig. 6C). Whereas arginine treatment delayed the resuscitation of persister bacteria because colony numbers in arginine-treated persisters were lower than in untreated persisters (Fig. 6D). Third, notably, the administration of exogenous L-serine and arginine also regulated resuscitation speed from meropenem-induced persister *E. coli* (Fig. S6A and S6B) and chloramphenicol-induced persister methicillin-resistant *Staphylococcus aureus* (MRSA) clinical isolates (Fig. S6C and D). This indicates that both metabolites function as widely conserved signals controlling persister resuscitation across Gram-negative and Gram-positive bacteria, including clinically isolated antibiotic-resistant pathogens. Therefore, we conclude that L-serine and arginine control the regrowth speed of antibiotic persister populations regardless of the specific bacterial species and antibiotics used.

**Figure 6.**
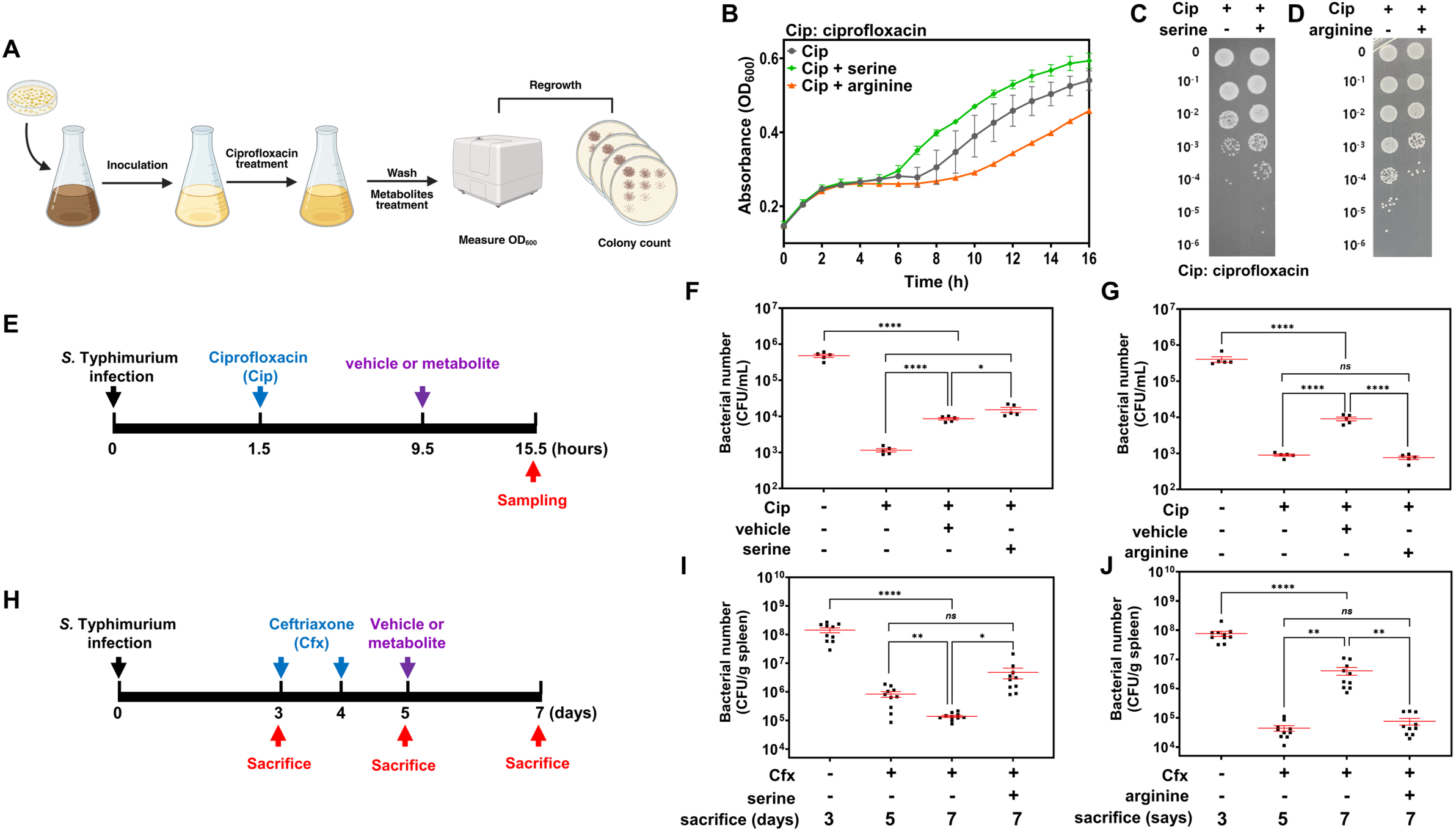
L-serine and arginine regulate the regrowth of antibiotic persisters within the host. (A–D) *In vitro* regrowth assay of antibiotic persister in laboratory medium. (A) Schematic of the *in vitro* regrowth assay workflow for antibiotic persister bacteria. Overnight cultures of *S*. Typhimurium ATCC 14028s were grown in N-minimal medium at 37 °C for 2.5 h with aeration until reaching the log phase. Ciprofloxacin (5 μg/mL) was then added for 3.5 h to induce persister formation. And then, samples were washed, and regrowth rate was monitored in fresh N-minimal medium with/without 20 mM L-serine or 28 mM arginine. Regrowth speed was measured by optical density (OD_600_) (B) and colony forming unit (CFU) enumeration (C and D). (B) Growth curves show that L-serine accelerates regrowth from antibiotic persistence, whereas arginine delays it compared to the untreated control. (C and D) CFU assay confirms that serine enhances regrowth from antibiotic persistence (C) while arginine suppresses regrowth of persister bacteria (D). (E–G) Regrowth assay of intracellular persister bacteria inside J774A.1 macrophage. (E) Schematic timeline of experimental observations regarding the survival of antibiotic persister bacteria within macrophages. J774A.1 macrophages were infected with *S*. Typhimurium at a multiplicity of infection (MOI) of 10 for 1.5 h, followed by treatment with ciprofloxacin (1 µg/mL) for 8 h to induce intracellular persister formation inside macrophages. After antibiotic removal, cells were incubated for 6 h in fresh medium containing vehicle (PBS) or metabolites (20 mM L-serine or 28 mM L-arginine). Total infection period was 15.5 h, and intracellular bacteria were enumerated by CFU counting at the indicated time points. (F) CFU counts indicate that L-serine significantly promotes resuscitation of intracellular persisters after antibiotic treatment. (G) Arginine treatment delays intracellular regrowth compared to the untreated control. (H–J) *In vivo* regrowth assay of antibiotic persister in murine models. (H) Schematic timeline of bacterial infection and treatment in mice. Overnight cultures of *S*. Typhimurium were harvested and suspended in PBS at 1.25 × 10 CFU/mL, and mice were intraperitoneally infected with 200 µL of the suspension (2.5 × 10³ CFU per mouse). Mice in the infection-only group were sacrificed at 3 days post-infection, whereas the remaining groups received intraperitoneal ceftriaxone (3 mg/mouse) on days 3 and 4 to induce antibiotic persister formation in mice. One day after the second antibiotic treatment (day 5), mice were intraperitoneally administered vehicle (0.85% saline) or L-serine or arginine (6 mg/mouse). Antibiotic-treated mice were sacrificed at day 5 (2 days after the first ceftriaxone dose) or day 7 (following metabolite or vehicle administration) for bacterial CFU enumeration in the spleen. (I) L-serine supplementation promotes the regrowth of antibiotic persister bacteria compared to the untreated group in mice models. (J) Arginine delays the resuscitation of antibiotic persister bacteria compared with the untreated group in mice. Data are presented as mean ± s.e.m. from biologically independent replicates. Statistical significance was determined using one-way ANOVA with Tukey’s multiple comparisons test; ns, not significant; \**P* < 0.05; \*\**P* < 0.01; \*\*\*\**P* < 0.0001.

### L-Serine and arginine regulate the regrowth speed of antibiotic persisters within macrophages and murine models during infection

Intracellular pathogens such as *Salmonella enterica* can infect and proliferate within macrophages(*38*). Although antibiotic therapy aims to eradicate intracellular *Salmonella*, persister populations persist within phagosomal vacuoles within macrophages, leading to recurrent infections and lethal systemic infections(*39*). Notably, *Salmonella* can readily generate a persister population inside the phagosome of macrophages, thereby being tolerant to antibiotics(*39*). In this study, we determined that L-serine and arginine can modulate the regrowth of antibiotic persisters (Fig. 6A-D). Thus, we hypothesized that L-serine and arginine may regulate recurrent *Salmonella* infections in the host.

To test this hypothesis, we infected macrophages with *Salmonella* and then administered antibiotics. Administration of ciprofloxacin, an antibiotic that is permeable to phagosomes, led to the retention of persister subpopulations within macrophage phagosomes (Fig. 6E-G). Subsequent to the elimination of antibiotics, we administered L-serine or arginine to assess the regrowth of persister bacteria within macrophages (Fig. 6E). Exogenous L-serine and arginine were administered at non-cytotoxic doses to macrophages (Fig. S7). Persister bacteria were revived more rapidly when exposed to L-serine within macrophages (Fig. 6F), but arginine treatment impeded the regrowth of persister bacteria (Fig. 6G). To ascertain if exogenous L-serine and arginine penetrate phagosomes (Fig. S8A), where persister bacteria are located, we quantified L-serine and arginine levels by mass spectrometry in *Salmonella*-containing vacuoles (SCV). L-serine and arginine effectively reached the phagosomal vacuoles after supplementation (Fig. S8B-C). Therefore, exogenous L-serine and arginine facilitate resuscitation within host cells during infections.

Additionally, we examined whether L-serine and arginine regulate the resuscitation speed of antibiotic persister bacteria in murine models. Mice infected with *Salmonella* were prompted to develop persister populations via ceftriaxone treatment, which penetrates phagosomes and is frequently employed to manage severe *Salmonella* infections in clinical environments(*40*) (Fig. 6H). Exogenous metabolites were provided 48 hours after antibiotic treatment (Fig. 6H). Ceftriaxone treatment successfully produces antibiotic persister populations in mice (Fig. 6I and H). Notably, the injection of L-serine resulted in a significantly increased resuscitation speed of antibiotic persister bacteria in mice compared to the untreated group (Fig. 6I). In addition, the addition of arginine to antibiotic persister bacteria in mice resulted in a significant delay in resuscitation speed compared to untreated samples (Fig. 6J). Collectively, these findings indicate that L-serine and arginine metabolites can serve as signals to regulate the resuscitation of antibiotic persister pathogens during infections in innate immune cells and in murine models.

## Discussion

Bacteria regulate their metabolism in response to environmental conditions, enabling pathogenic bacteria to cause diseases in the host(*41*). Bacteria can enter a dormant state under unfavorable conditions, allowing them to become antibiotic persister bacteria that tolerate immune responses and antibiotics, thereby facilitating the establishment of chronic infections(*17*). This study demonstrates that dormant bacteria selectively regulate their metabolic pathways to enable swift resuscitation from dormancy when conditions become favorable again (Fig. 7). We performed metabolome and proteome profiling analysis, along with in silico modeling, to identify specific metabolic pathways that are differentially regulated during dormancy to promote resuscitation speed (Fig.1 and Figs. S1 and S2). L-serine promotes the regrowth of dormant bacteria (Fig. 2). In contrast, arginine inhibits the regrowth of dormant bacteria from their dormant state (Fig. 4). Furthermore, we demonstrate that L-serine and arginine regulate specific pathways during regrowth that govern the resuscitation of dormant bacteria (Fig. 3 and 5). L-serine and arginine also regulate resuscitation speed of antibiotic persister bacteria, which is representative dormant bacteria (Fig. 6). These characteristics are validated in both in vitro cell culture models and in vivo mouse studies (Fig. 6). Notably, to our knowledge, for the first time, we identify specific metabolites to regulate the regrowth speed of antibiotic persister during infections (Fig. 6). This research represents a paradigm shift, as it has been established that dormant bacteria selectively regulate specific metabolic pathways, rather than overall metabolism, which in turn influences the speed of regrowth from dormancy during infection (Fig. 7).

**Figure 7.**
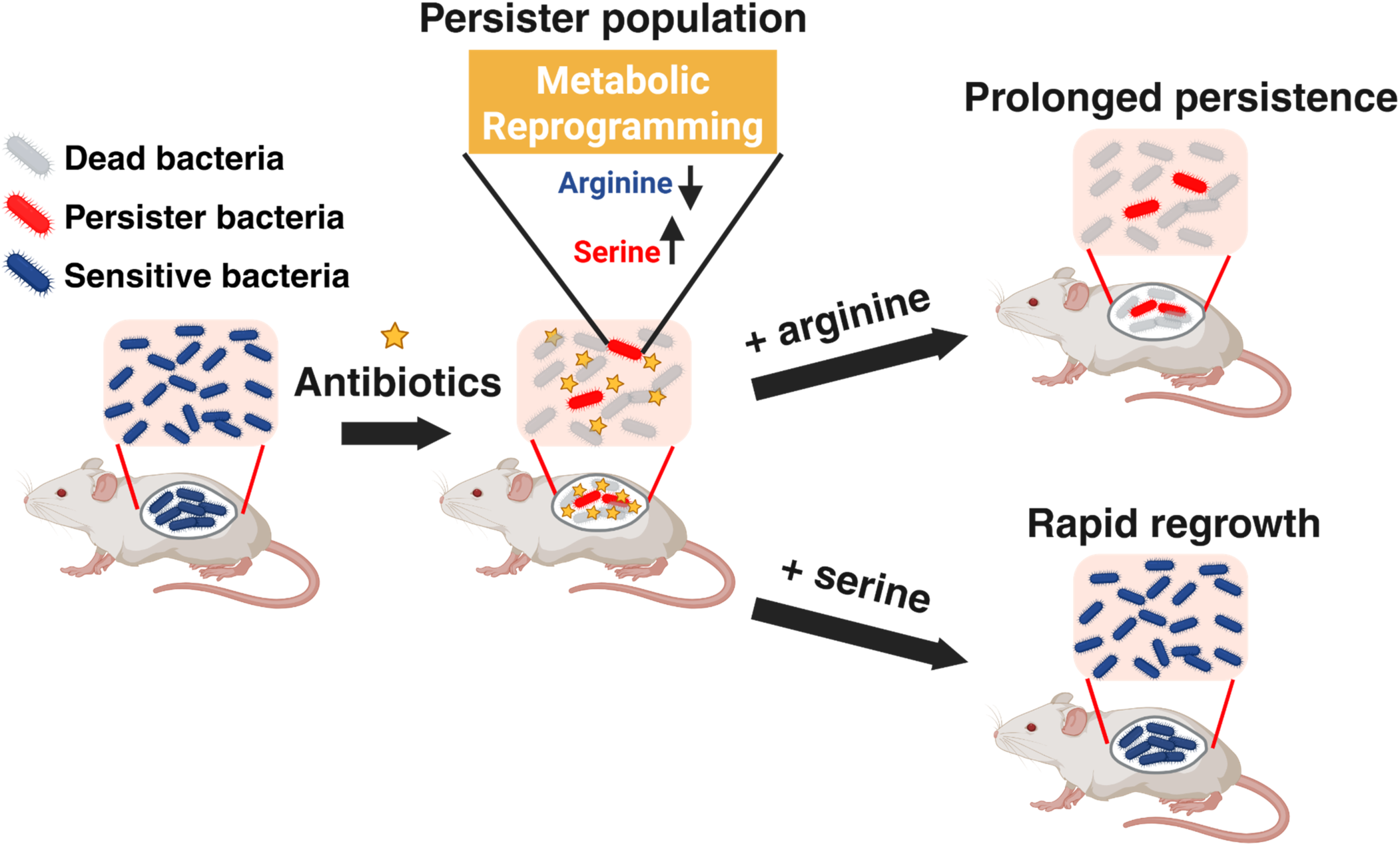
Metabolic signals control the resuscitation speed of antibiotic persister pathogens during infection. A schematic illustration suggests that metabolic signals, L-serine and arginine, regulate the regrowth speed of antibiotic persister bacteria in the host. Persister bacteria selectively reprogram their metabolism by accumulating L-serine while reducing arginine levels to promote rapid wake-up from dormancy. Given that antibiotic persistence is one of the reasons for bacterial chronic infections in clinical settings, and that exogenous metabolite treatment modulates persister regrowth dynamics *in vivo*, it provides a potential new approach to control chronic infection.

L-serine is highly accumulated in fast-regrowing dormant bacteria (Fig. 1). During infections, L-serine increases growth fitness in enteric bacterial species, advantaging pathogenic bacteria against competitors(*42*), and serves as a carbon source under starved conditions like urine(*43*). L-serine also inhibits dormant spore production in *Clostridium*, reducing dormancy by inhibiting early spore formation and stimulating continuous growth(*44*), thereby consistently facilitating escape from dormancy in this study (Fig. 2). Additionally, L-serine fluxes into glutathione metabolism via glycine, which is critical for oxidative stress defense(*45*). Since cysteine metabolism, required for oxidative stress defense, increases highly with L-serine treatment in dormant bacteria (Fig. 5C), L-serine accumulation during dormancy may contribute to oxidative stress defense. Increased glutathione and cysteine synthesis decrease the level of reactive oxygen species (ROS)(*46*), which innate immune cells use to inhibit bacterial growth. Notably, we determined that L-serine is a signal, not a metabolic precursor, because an L-serine analog, which bacteria can’t degrade and use as a carbon source, also activates the resuscitation from dormancy. Thus, the signal molecule L-serine stimulates the rapid regrowth of dormant bacteria and antibiotic persister cells by promoting metabolic reprogramming under unfavorable conditions.

Arginine plays a critical role in biofilm formation and the regulation of metabolic pathways in several pathogenic bacteria(*47–49*). Supplementation with arginine inhibits glycolysis and peptidoglycan synthesis in *Streptococcus*(*50*). Fast-regrowing dormant bacteria exhibit elevated activity in the glycolysis pathway (Fig. 1), which may facilitate cell wall synthesis by producing peptidoglycan precursors during regrowth. Consequently, low arginine levels in dormant bacteria are essential for maintaining high glycolytic activity, enabling swift resuscitation. Arginine also facilitates biofilm formation in Enterococcus faecalis(*49*). During biofilm formation, the bacterial growth rate declines while dormant cell populations rise(*51*), because biofilms comprise heterogeneous populations including dormant subpopulations(*52*). Thus, bacteria can enhance dormant populations by forming biofilms in high arginine environments, aligning with our findings that arginine inhibits bacterial regrowth from dormancy and preserves dormant status (Fig. 4). In addition, ppGpp is an essential molecule for adaptation to harsh environments, facilitating the production of dormant bacteria by reducing ribosome synthesis(*53*) and activating stress response systems during dormant states(*54*). The supplementation of arginine reduces ppGpp levels(*55*), leading to the dormant bacteria’s inability to adapt to unfavorable conditions during dormancy. Furthermore, arginine supplementation inhibits fumarate synthesis, which can facilitate rapid regrowth by functioning as a cofactor for the transcriptional regulator ArgR(*56*) (Fig. 4I). Therefore, bacteria can prolong a dormant state by accumulating arginine, a common pathway for extending dormancy.

Antibiotic persistence contributes to chronic infections by enhancing tolerance to antibiotic treatment in clinical environments(*57, 58*). During antibiotic treatment, pathogenic bacteria enter a dormant persister state and reactivate once conditions become favorable. For several decades, numerous studies on antibiotic persistence have been conducted in laboratory settings(*59*). But recent research suggests that diverse nutrient sources and DNA repair mechanisms are essential for the regrowth of dormant bacteria within hosts(*60, 61*). In this study, we demonstrate that certain metabolite signals regulate the resuscitation speed of antibiotic persisters in innate immune cells and mice during infection (Fig. 6). This indicates the possibility of modulating the regrowth duration of antibiotic persisters through metabolite treatment, thereby establishing a new strategy for managing antibiotic persister pathogens and chronic infections (Fig. 7). Especially, this study provides insights for developing therapeutics targeting persisters, since regrowing dormant bacteria are sensitive to antibiotics(*62*), potentially allowing us to determine when persister bacteria wake from dormancy and can be effectively killed during infection. Notably, the mechanisms are applicable to clinically isolated MRSA and *E. coli* (Fig. S6). Thus, the metabolic signal-mediated dormancy exit mechanism is not restricted to model organisms and is widely conserved across bacteria. Moreover, the conservation of the dormant phenotype in diverse pathogenic bacteria(*2*) and drug-resistant cancer cells(*63*) suggests that signal molecules regulating dormancy and regrowth may be broadly applicable for controlling various dormant cell types.

## Acknowledgments

We are grateful to all of authors and Yeom lab members for comments on the early draft of manuscript. This work was supported by the New Faculty Startup Fund from Seoul National University to J.Y., the National Research Foundation Korea Basic Science Research Programs (2021R1C1C1005184, 2020M3A9H5104237 and 2020R1A5A1019023 to J.Y., RS-2024-00345936 to G.H.), Creative-Pioneering Researchers Program through Seoul National University, the Korea Health Technology R&D Project through the Korea Health Industry Development Institute (KHIDI) (grant number: HI23C026400 and RS-2023-00304637), the Global-LAMP Program of the National Research Foundation of Korea (NRF) grant funded by the Ministry of Education (No. RS-2023-00301976) and), the Korea Basic Science Institute (No. C523400 to G.H.).

## Contributions

J.M.H., B.S. and J.Y designed the study. B.S., D.H. and S.N.L. carried out the proteome analysis. J.L., J.L., E.C. and G.S.H carried out metabolome analyses. J.K., D.K. Y.B. and K.K. performed animal experiments. Y.Q.L., L.K. and D.Y.L. conducted *in silico* analysis. D.L., K. J. and K.K. analyzed incorporated proteome and metabolome data. T.Y. and M.K. conducted experiments for molecular mechanism and regrowth. J.M.H, T.Y. and J.Y mainly wrote the paper with contributions from all authors.

## Supplementary Materials for

### The PDF file includes

Materials and Methods

Figs. S1 to S8

Tables S1 to S2

## Materials and Methods

### Microbial strains, plasmids, and growth conditions

Bacterial strains and plasmids used in this study are presented in Supplementary Table S1. All *S.* Typhimurium strains are derived from strain wild-type strain 14028s. All *E. coli* strains are derived from strain wild-type strain MG1655 K12. Methicillin-resistance *staphylococcus aureus* (MRSA) are obtained from Seoul National University Bundang Hospital. DNA oligonucleotides used in this study are presented in Supplementary Table S2. *S.* Typhimurium were grown at 37 °C in Luria-Bertani broth (LB) and N-minimal media (pH 7.7) supplemented with 0.1% casamino acids, 38 mM glycerol, and concentrations of 1mM MgCl_2_. *E. coli* and MRSA were grown at 37 °C in M9 minimal media (pH 7.7) containing 1 x M9 salts, 2 mM MgSO_4_, 0.1 mM CaCl_2_, 1% glucose, 1% casamino acids, 1 mM Thiamine-HCl and 0.05 mM nicotinamide.

### Construction of bacterial mutants

Bacterial chromosomal mutants were constructed using the one-step disruption method with minor modifications. To construct the *serA* (TY40) and *argH* (TY37) mutants, a kan cassette was introduced into the *serA* and *argH* gene as follows: A kat gene fragment was amplified from pKD4 using primers B15H8/B15H6 (*serA)*, B15F9/B15G1 (*argH)* and then introduced into wild-type *S*. Typhimurium (14028s) harboring plasmid pKD46 selecting from kanamycin resistance. The resulting strain (TY36 and TY32) was kept at 30 °C and transformed with pCP20 to remove the cat cassette. To construct strains specifying a C-terminally HA-tagged SdaA protein (TY44), SerA protein (TY45) and ArgH protein (TY46), a kan cassette was introduced at the 3′ end of the *serA*, *sdaA* and *argH* gene: A kan gene fragment was amplified using primers B20A5/B20A6 for TY44, B20A3/B20A4 for TY45 and B20A7/B20A8 for TY46 from plasmid pKD4. The amplified fragments were introduced into wild-type *S*. Typhimurium (14028s) harboring plasmid pKD46 selecting for kanamycin resistance.

A plasmid expressing SdaA and ArgH was constructed as follows: the *sdaA* and *argH* gene was amplified using primers B14I7/B14I8 for *sdaA,* B15F9/B15G1 for *argH* and then were introduced between the *BamHI* and *HindIII* sites of plasmid pUHE-21-2-*lacI^q^*.

### Western blot assay

Bacteria cells were grown in N-minimal medium. Each 16 hours overnight cultured bacteria strains are inoculated in N-minimal medium with 1/100 dilution and treated with chloramphenicol (20 `µ`g/mL) at 3 h after inoculation. At specific time point, crude extracts were prepared in B-PER reagent (Pierce) with 100 `µ`g/mL lysozyme and EDTA-free protease inhibitor (Roche). Samples were loaded on 4–12% Bolt Bis-Tris gels (Life Technologies) then transferred to nitrocellulose membrane using the iBlot2 machine (Life Technologies). Membranes were blocked with 5% skim milk solution (bovine serum albumin (BSA) in Tris-buffered saline with 0.1% Tween-20 (TBST) at room temperature (RT) for 2 h. Then, samples were analyzed using antibodies against the HA peptide or the AcnB or GroEL proteins. Rabbit anti-AcnB (LifeSpan BioSciences, LS-C486267), anti-HA (Sigma-Aldrich, H6908) antibodies were used at 1:2,000 dilution, and anti-groEL (Abcam, ab82592) were used at 1:5,000 dilution. Secondary horseradish peroxidase-conjugated anti-rabbit (Cytiva, NA934) were used at 1:5,000 dilution. After washing with TBST 3 times, blots were developed with the ECL Western Blotting Substrate (Pierce), detected by Amersham Imager AI680(GE healthcare).

### Measurement of ATP amounts

Each 16 hours overnight cultured bacteria strains were inoculated in N-minimal medium with 1/100 dilution and treated with chloramphenicol (20 `µ`g/mL) or tetracycline (4 `µ`g/mL) at 3 h after inoculation. 3 h after antibiotic treatment bacterial samples are inactivated by 70 °C heating for 10 min. Intracellular ATP was measured by microplate reader (Biotek, Synergy H1) using the BacTiter-Glo Microbial Cell Viability Assay Kit (Promega) according to the manufacturers’ instructions. ATP measurements were normalized by optical density at 600 nm (OD_600_).

### Regrowth Assay

Overnight cultured bacteria strains were inoculated in N-minimal or M9 medium with 1/100 dilution and treated with chloramphenicol (20 `µ`g/mL or 50 `µ`g/mL), tetracycline (4 `µ`g/mL), ampicillin (30 `µ`g/mL), ceftriaxone (30 `µ`g/mL), meropenem (0.75 µg/ml), or ciprofloxacin (1.6 `µ`g/mL or 5 `µ`g/mL) at 3 h after inoculation. 3 h after antibiotic treatment cells were analyzed by optical density at OD_600_ and washed with fresh N-minimal media twice.

For measuring the OD_600_ to investigate escape from the slow-growth state into the growth state, 1/100 diluted samples were inoculated into fresh N-minimal or M9 medium with or without metabolites (L-serine 30 mM, alpha-methyl-L-serine 30 mM, or arginine 28 mM). Microbial growth was measured using by microplate reader (Biotek, Synergy H1) at 37 °C for 16 h.

To count the colony forming unit (CFU) of the persister bacteria, antibiotic treated samples were washed and re-inoculated into fresh N-medium with or without metabolites (L-serine 30 mM or arginine 28 mM) and incubated in test tubes until 24 h. At 6 h after incubation, cells were harvested and measured by optical density at 600nm. Samples were washed with N-minimal medium twice. After serial dilution, 5`µ`L of each samples were spotted on the Luria-Bertani (LB) plates.

### Quantitative RT-PCR

To measure mRNA abundance, cells were grown in N-minimal medium containing 1mM MgCl_2_ in 38 mM of glycerol at 37 °C for 6 h. Total RNA was purified by using RNeasy Kit (Qiagen) with on-column DNase treatment, and complementary DNA was synthesized using VILO Super Mix (Life Technologies). Quantification of transcripts was carried out by qRT-PCR using SYBR Green PCR Master Mix (Applied Biosystems) in a QuantStudio 6 Flex Real-Time PCR System (Applied Biosystems). mRNA abundance was determined by using a standard curve obtained from PCR products generated with serially diluted genomic DNA, and results were normalized to the levels of the ompA gene. Data shown are an average from at least three independent experiments. Primers used in the qRT-PCR assay are presented in Supplementary Table S2.

### Intramacrophage Regrowth Assay

The murine-derived macrophage cell line J774A.1 was cultured in Dulbecco’s modified Eagle’s medium (DMEM; Welgene) supplemented with 10% fetal bovine serum (FBS, Life Technologies) at 37 °C under 5% CO_2_. Confluent monolayers were prepared in 24-well tissue culture plates. Each well of a 24-well plate was seeded with 5 × 10^5^ cells suspended in DMEM/10% FBS and incubated at 37 °C under 5% CO_2_ for 20 h. Bacteria were grown in LB media at 37 °C for 16 h. Bacterial cells were washed two times with phosphate-buffered saline (PBS, Welgene), suspended in pre-warmed DMEM, and then added to the cell monolayer at a multiplicity of infection (MOI) of 10. Following 30 min incubation, the wells were washed three times with pre-warmed Dulbecco’s PBS (DPBS, Welgene) to get rid of extracellular bacteria and then incubated with pre-warmed medium supplemented with 120 μg/ml gentamicin for 1 h to kill the remaining extracellular bacteria. Then, the wells were washed three times with DPBS and incubated with pre-warmed medium supplemented with 10 μg/ml gentamicin and 1 μg/ml ciprofloxacin. After 8 h incubation, wells were washed three times with DPBS and incubated with pre-warmed medium supplemented with 10 μg/ml gentamicin and metabolites (L-serine 20 mM or arginine 28 mM) and incubated for 6 h. Wells were washed two times with DPBS to remove extracellular bacteria. Then, 1 ml DPBS and 0.1% Triton X-100 was added to the wells, and following a 10 min incubation, serial dilutions of bacteria were plated on LB agar plates and incubated at 37 °C O/N to determine the number of colony forming unit (CFU).

### MS analysis for proteome profiling in dormant and regrowing bacteria

Proteome analysis of dormant and regrowing bacteria was conducted by Protein and Proteomics Centre in National University of Singapore (NUS PPC). We employed the SWATH-MS technique to examine the global proteome in latent and proliferating bacteria (Fig. 1A-B) with slight modifications as previous methods (*64*).

We employed in-solution digestion and desalting for protein extraction. Prior to digestion, protein concentration was determined using a BCA protein assay kit. For each sample, 50 µg of protein was transferred into a microcentrifuge tube. The samples were then reduced by adding dithiothreitol (DTT) to a final concentration of 10 mM and incubating at 56°C for 30 minutes. Following reduction, iodoacetamide (IAA) was added to a final concentration of 55 mM and incubated in the dark at RT for 30 minutes to alkylate the cysteine residues. The samples were then diluted with 50 mM ammonium bicarbonate to a final concentration of 1 M urea or less. Trypsin was added at a protein-to-enzyme ratio of 50:1 (w/w), and the mixture was incubated at 37°C overnight (16–18 hours). The reaction was finally quenched by adding formic acid to a final concentration of 1%.

The digested peptide solution was desalted to remove salts and detergents that could interfere with mass spectrometry analysis. For this process, C_18_ solid-phase extraction (SPE) cartridges or tips were used. The cartridges were first conditioned with 100% methanol or acetonitrile, followed by an equilibration with 0.1% formic acid in water. The acidified peptide solution was then loaded onto the conditioned cartridges. To remove salts and contaminants, the cartridges were washed with 0.1% formic acid in water. The peptides were subsequently eluted with a solution of 80% acetonitrile and 0.1% formic acid. Finally, the eluted peptides were dried using a speed vacuum concentrator and resuspended in 0.1% formic acid in water for subsequent LC-MS analysis.

Subsequently, the desalted peptide samples were analyzed using a liquid chromatography system coupled to a mass spectrometer. All 24 samples will be analyzed individually without chemical labeling, employing a label-free data-independent acquisition (DIA) quantitative methodology known as SWATH. For SWATH-MS analysis, we employed Eksigent nanoLC reversed phase chromatography for liquid chromatography. Mass analysis was conducted using the SCIEX TripleTOF 6600 MS in long gradient mode, incorporating suitable blank runs. Subsequently, we constructed a spectrum library and examined SWATH to identify proteins from the samples.

### Sample preparation for MS analysis for proteome profiling by exogenous treatment of metabolites in dormant bacteria

Bacterial pellets were resuspended in lysis buffer (4% SDS, 100 mM Tris-HCl; pH 7.5, 100 mM Dithiothreitol [DTT]) and lysed using a probe sonicator. The lysates were heated at 95 °C for 15 min. After cell debris was removed by centrifugation at 15,000 rpm for 10 min at RT, protein concentration was determined using a reducing agent-compatible BCA assay kit (Thermo Fisher Scientific, Waltham, MA, USA).

For protein digestion, 100 µg of protein was precipitated using cold acetone at a 1:6 (v/v) ratio and incubated overnight at –20 °C. The protein pellets were resuspended in denaturation buffer (2% SDS, 100 mM Tris-HCl; pH 8.5, 10 mM tris(2-carboxyethyl)phosphine [TCEP], 50 mM 2-Chloroacetamide [CAA]). Protein digestion was performed using the filter-aided sample preparation (FASP) method, as described previously (*65, 66*). The denaturated and alkylated protein sample was loaded onto a 30 kDa Amicon filter (Merck Millipore, Darmstadt, Germany) and centrifuged at 14,000 × g for 15 min, then diluted with 400 µL of urea buffer (8 M Urea, 100 mM Tris-HCl; pH 8.5, 0.2 µm filtered). This step was repeated three times, followed by three washes with 50 mM HEPES buffer (pH 8.5). The proteins were digested overnight at 37 °C with a trypsin/Lys-C mixture (enzyme-to-protein ratio, 1: 100 w/w). After digestion, peptides were collected by centrifugation into a new Eppendorf tube. Peptide concentrations were measured by tryptophan fluorescence assay (*67*).

The peptides were labeled with a 16-plex TMTpro isobaric label reagent (Thermo Fisher Scientific, Waltham, MA, USA) following the manufacturer’s instruction at a sample-to-tag ratio of 1:5 (w/w). A total of 24 samples were analyzed using two TMTpro 16-plex sets. For each sample, 10 µg of peptide was used for labeling, and 10 µL of TMTpro reagent solution was added along with acetonitrile (ACN). To minimize potential batch effects, the samples were randomly assigned to channels. The labeled peptides were then desalted using a Sep-Pak tC18 cartridge (Waters, Milford, MA, USA). Desalted peptides were fractionated using an Agilent 1290 HPLC system (Agilent, Santa Clara, CA, USA) for 55 min run method as previously mentioned (*68*). Overall, 96 fractions were concatenated into 24 fractions. The fractionated samples were dried using a Speed-Vac and stored at –80 °C until further analysis.

### LC-MS/MS analysis for proteome profiling by exogenous treatment of metabolites in dormant bacteria

Fractionated peptides were resuspended in Solvent A (0.1% formic acid in HPLC water) for MS analysis and analyzed using an Ultimate 3000 RSLC System (Dionex, Sunnyvale, CA, USA) coupled with an Orbitrap Exploris 480 mass spectrometer (Thermo fisher scientific, Waltham, MA, USA). Each sample was loaded onto a trap column (Acclaim PepMap100-C18, 3 µm, 75 µm x 2 cm, Thermo Fisher Scientific, Waltham, MA, USA) and the peptides were separated on an analytical column (IonOpticks Aurora C18, 75 µm x 60 cm, 1.7 µm). The samples were analyzed using a 150 min gradient from 5% to 95% Solvent B (0.1% formic acid in acetonitrile) at 200 nL/min. MS1 scans were acquired in the range of 350-1800 m/z with a resolution of 120,000 at 200 m/z. MS/MS analyses were performed using high-energy collisional dissociation (HCD) with a normalized collision energy of 32 and a resolution of 30,000 at 200 m/z. Data-dependent acquisition was used to select the top 15 most abundant precursor ions, with an isolating window of 0.7 m/z.

### Protein identification for proteome profiling by exogenous treatment of metabolites in dormant bacteria

MS spectra files were processed using Fragpipe (v22.0) (*69*) against the Uniprot *Salmonella* typhimurium ATCC 14028 database, containing 5,376 entries plus common contaminants. Trypsin and Lys-C were used as the digestion enzymes for the identification of tryptic peptides. Fully tryptic peptides with up to a maximum of two missed cleavages and lengths of 6-50 amino acids were allowed. The precursor and fragment mass tolerance were set to 20 ppm and 0.02 Da, respectively. Variable modifications included methionine oxidation (+15.995 Da) and N-terminal acetylation (+42.01056 Da). TMTpro 16plex N-term and lysine residues (+304.20715 Da) and carbamidomethylation of cysteine (+57.02146 Da) were set as fixed modifications. Peptide and peptide spectrum matches (PSMs) were validated using percolator based on a 1% false discovery rate (FDR).

### Statistical Analysis for proteome profiling by exogenous treatment of metabolites in dormant bacteria

Statistical analysis of TMT data was performed using Perseus software (*70*). Log2 transformation followed by width-adjusted normalization was applied for pairwise quantification. To identify differentially expressed proteins (DEPs), Student’s t-test was performed using p-value threshold of 0.05 and a fold-change cutoff of 1.5. Proteins upregulated at the regrowth early point were further classified based on their expression at the middle point. The proteins that remained upregulated were defined as sustained, while those not differentially expressed at the middle point were considered early point-specific. Gene Ontology (GO) functional annotation analysis of the eight groups was conducted using database for annotation, visualization and integrated discovery (DAVID) (p-value < 0.05, count ≥ 3). For each group, the three GO terms with the most significant p-values were selected for further analysis. Hierarchical clustering of proteins annotated with these GO terms was performed based on Z-score normalized intensities, and the heatmap was visualized using Python 3.11.8 (Seaborn (v.0.13.2), Matplotlib(v.3.8.3), SciPy(v.1.12.0), Pandas(v.2.2.1).

### Metabolomic profile analysis

To extract intracellular metabolites of *Salmonella* Typhimurium, we used modified method as described previously (*71*). For quenching cellular metabolism, 5 mL of the cell culture (OD₆₀₀ = 0.8) was rapidly snap-frozen in liquid nitrogen for 1 min, followed by thawing on ice. Samples were centrifuged at 12,000 × *g* for 5 min at –10 °C to pellet the biomass. The supernatant was carefully removed, and the resulting cell pellets were either subjected immediately to extraction or stored at –80 °C until further processing. Cell pellets were resuspended in 2 mL of pre-chilled (–40 °C) 100% methanol. The suspension was snap-frozen in liquid nitrogen for 8 min, thawed on wet ice, vortexed for 30 s, and the freeze–thaw cycle was repeated twice more. Following extraction, samples were centrifuged at 14,800 × *g* for 8 min at –10 °C, and the supernatant was transferred into fresh 2 mL microcentrifuge tubes. Extract volumes were normalized according to the initial OD₆₀₀ value of each culture by adjusting with the extraction solvent. Quality control (QC) samples were prepared by pooling equal aliquots from all samples. The extract was dried under vacuum and stored at –80 °C until analysis. Immediately prior to LC–MS analysis, dried extracts were reconstituted in 300 μL of a water-acetonitrile mixture (1:3, v/v) containing an internal standard mixture. The internal standard mixture consisted of isotope-labeled metabolites, including betaine-d₁₁ (0.1 μg/mL), glutamic acid-¹³C₅ (10 μg/mL), leucine-¹³C₆ (5 μg/mL), uridine-¹³C₉,¹⁵N₂ (10 μg/mL), phenylalanine-¹³C₆ (2 μg/mL), succinic acid-¹³C₄ (10 μg/mL), and taurine-¹³C₂ (10 μg/mL).

Intracellular metabolite profiling was carried out on an ACQUITY™ ultra-performance liquid chromatography (UPLC) system (Waters, Manchester, UK) coupled to a TripleTOF® 5600 quadrupole time-of-flight mass spectrometer (QTOF MS, Sciex, Concord, Canada). Chromatographic separation was achieved using a SeQuant ZIC-HILIC column (2.1 × 100 mm, 3.5 `µ`m, 200 Å; Merck, Darmstadt, Germany) maintained at 35 °C with a flow rate of 0.4 mL/min. The mobile phase consisted of (A) 0.1% formic acid and 10 mM ammonium acetate in acetonitrile–water (9:1, v/v) and (B) 0.1% formic acid and 10 mM ammonium acetate in acetonitrile–water (1:1, v/v). The UPLC gradient was programmed as follows: 1% B for 2 min, ramped to 55% B from 2 to 8 min, further increased to 99% B from 8 to 9 min, held for 2 min, then decreased to 1% B from 11 to 11.1 min, and maintained at 1% B for 3.9 min to equilibrate the column prior to the next run.

A 5 `µ`L aliquot of each sample was injected and analyzed in both positive and negative electrospray ionization (ESI) modes over a mass range of *m/z* 30–1,000. The ion source parameters were as follows: curtain gas, 30 psi; nebulizer gas, 60 psi; heater gas, 60 psi; source temperature, 500 °C; and ion spray voltage, +5,500 V in positive mode and −4,500 V in negative mode. Mass accuracy was maintained using an atmospheric pressure chemical ionization (APCI) calibration solution delivered via an automated calibrant delivery system (Sciex, Concord, Canada) integrated with a DuoSpray™ ion source.

To ensure data consistency, quality control (QC) samples were analyzed following every six analytical sample injections. The coefficient of variation (CV) for QC samples was determined as an indicator of measurement stability. UPLC/QTOF-MS datasets were processed using MarkerView software (v1.3.1, Sciex, Concord, Canada). Spectral features were extracted, aligned, and compiled into peak tables containing *m/z* values and retention times for each sample. Data were normalized to the total ion intensity. Identification of metabolites with CV values less than 20% were performed by comparing accurate mass measurements, fragment ion spectra, and retention times against the Human Metabolome Database (HMDB), METLIN, and an in-house reference library.

### Flux Balance Analysis and Flux Visualization

Flux simulations were performed using the genome-scale *Escherichia coli* model *i*ML1515 within the COBRA Toolbox (*72*) in MATLAB R2022b and the Gurobi (http://www.gurobi.com) optimization solver. To mimic nutrient-limited, starvation-like conditions, uptake fluxes for all components of M9 minimal medium were restricted to 3% of their original composition. To model energy homeostasis under minimal growth, the biomass reaction was constrained to a fixed growth rate of 0.01 h⁻¹. The cellular objective was then set to maximization of ATP synthesis. The non-growth–associated ATP maintenance reaction (ATPM) was disabled by setting its lower bound to zero, allowing ATP turnover to be determined only by network demands under the specified growth and nutrient constraints. Next, flux balance analysis with molecular crowding (FBAwMC) was used to compute feasible steady-state flux distributions (*73*). A total of 5,000 flux solutions were generated to capture the variability of alternate metabolic states consistent with the imposed constraints.

To identify metabolic states with the highest and lowest ATP availability, percentile-based tail binning was applied to ATP flux values across all flux solutions. The ATP production capacity of each solution was quantified using flux-sum analysis of ATP-producing and ATP-consuming reactions, following established protocols (*74*). Flux solutions within the top 5% and bottom 5% of the ATP flux-sum distribution were designated as the “high-ATP” and “low-ATP” groups, respectively. For each group, the flux values of all reactions were averaged across individual solutions to obtain a representative metabolic flux distribution. The group-averaged fluxes were then mapped onto the corresponding reactions in the metabolic network and visualized using Escher (*75*).

### Determination of L-serine, arginine, and fumarate from vacuoles and bacteria

Metabolites, including L-serine, arginine and fumarate, from *Salmonella* Typhimurium and the SCV were extracted using the same method as in the above metabolomic analysis. To determine the levels and concentrations of L-serine, arginine and fumarate in *Salmonella* Typhimurium and SCV, an Agilent 1290 Infinity UPLC and an Agilent 6495 Triple Quadrupole MS system equipped with Agilent Jet Stream ESI source (Agilent Technologies, USA). Data acquisition and processing were performed using MassHunter Workstation software (version B.06.00; Agilent Technologies, USA). Chromatographic separation was achieved using a Waters Acuquity UPLC BEH Amide column (100 × 2.1 mm, 1.7 `µ`m; Waters, Manchester, UK) for L-serine and arginine, and Acclaim™ Organic Acid column (150 mm x 2.1 mm, 3 `µ`m; Thermo Scientific™) for fumarate. For L-serine and arginine analysis, the column temperature was maintained at 30 °C with a flow rate of 0.4 mL/min. The mobile phases consisted of 0.2% formic acid in water (A) and 0.2% formic acid in acetonitrile (B). The gradient program was as follows: 90% B for 2.0 min, 90–48% B over 5.0 min, 48% B for 2.0 min, 48–90% B over 1.0 min, and 90% B for 3.0 min, giving a total run time of 13 min.

For fumarate analysis, the column temperature was set to 40 °C with a flow rate of 0.3 mL/min. The mobile phases were 0.2% formic acid in water (A) and 0.2% formic acid in methanol (B). The gradient program was: 0% B for 1.0 min, 0–50% B over 7.0 min, 50–100% B over 2.0 min, 100% B for 2.0 min, 100–0% B over 0.1 min, and 0% B for 2.9 min, for a total run time of 15 min.

MS/MS analyses were performed in positive ion mode for L-serine and arginine, and in negative ion mode for fumarate, using the following parameters: capillary voltage, 3.5 kV; nebulizer nitrogen pressure, 30 psi; drying gas temperature, 200 °C; drying gas flow rate, 16 L/min; sheath gas temperature, 300 °C; sheath gas flow rate, 11 L/min; and nozzle voltage, 1500 V. Detection was performed using optimized multiple reaction monitoring (MRM) transitions as follows: L-serine (*m/z* 106 > 60; *m/z* 106 > 42), L-serine-¹³C₃ (*m/z* 109 > 91.1; *m/z* 109 > 62.1), arginine (*m/z* 175.1 > 115.9; *m/z* 175.1 > 70.3), arginine-¹³C₆ (*m/z* 181.2 > 121; *m/z* 181.2 > 74), fumarate (*m/z* 115 > 115.2), and fumarate-¹³C₂ (*m/z* 117 > 72).

### *In vivo* infection of murine models

Animal experiments were approved by the Institutional Animal Care and Use Committee (IACUC) of Seoul National University (SNU-220722-1-2) and University of Illinois at Chicago. Female C57BL/6 mice (7–8 weeks old) were obtained from Orientbio (Seongnam, Republic of Korea) and Jackson Laboratory (USA). For each experimental set, mice were randomly assigned to one of four treatment groups: an infection-only group, a ceftriaxone-treated group, and a regrowth phase with/without an amino acid.

*S.* Typhimurium ATCC 14028s grown overnight was harvested and suspended in PBS at 1.25 × 10⁴ CFU/ml. The mice were intraperitoneally infected with 200 µl of the bacterial suspension. Mice in the infection-only group were sacrificed at 3 days post-infection, while the remaining groups received intraperitoneal ceftriaxone (3 mg/mouse) at days 3 and 4 post-infection. One day after the second antibiotic dose, mice were intraperitoneally administered either 0.85% saline or amino acids (6 mg/mouse). Antibiotic-treated mice were then sacrificed at 5 days post-infection (i.e., 2 days after the initial antibiotic treatment) or at 7 days post-infection following metabolite administration.

Spleens were harvested upon sacrifice, weighed, and homogenized in 1 ml of ice-cold PBS. Homogenates were centrifuged at 300 × g for 3 min at 4°C to remove mammalian cells. The resulting supernatants were centrifuged at 5,000 × g for 5 min at 4°C, and the bacterial pellets were resuspended in cold PBS, serially diluted, and plated on LB agar to enumerate CFU.

### Quantification and statistical analysis

Data and statistical analysis were performed using GraphPad Prism 10.2.2. Replicates and statistical details can also be found in the methods and figure legends.

## Supplemental figures

**Fig. S1.**
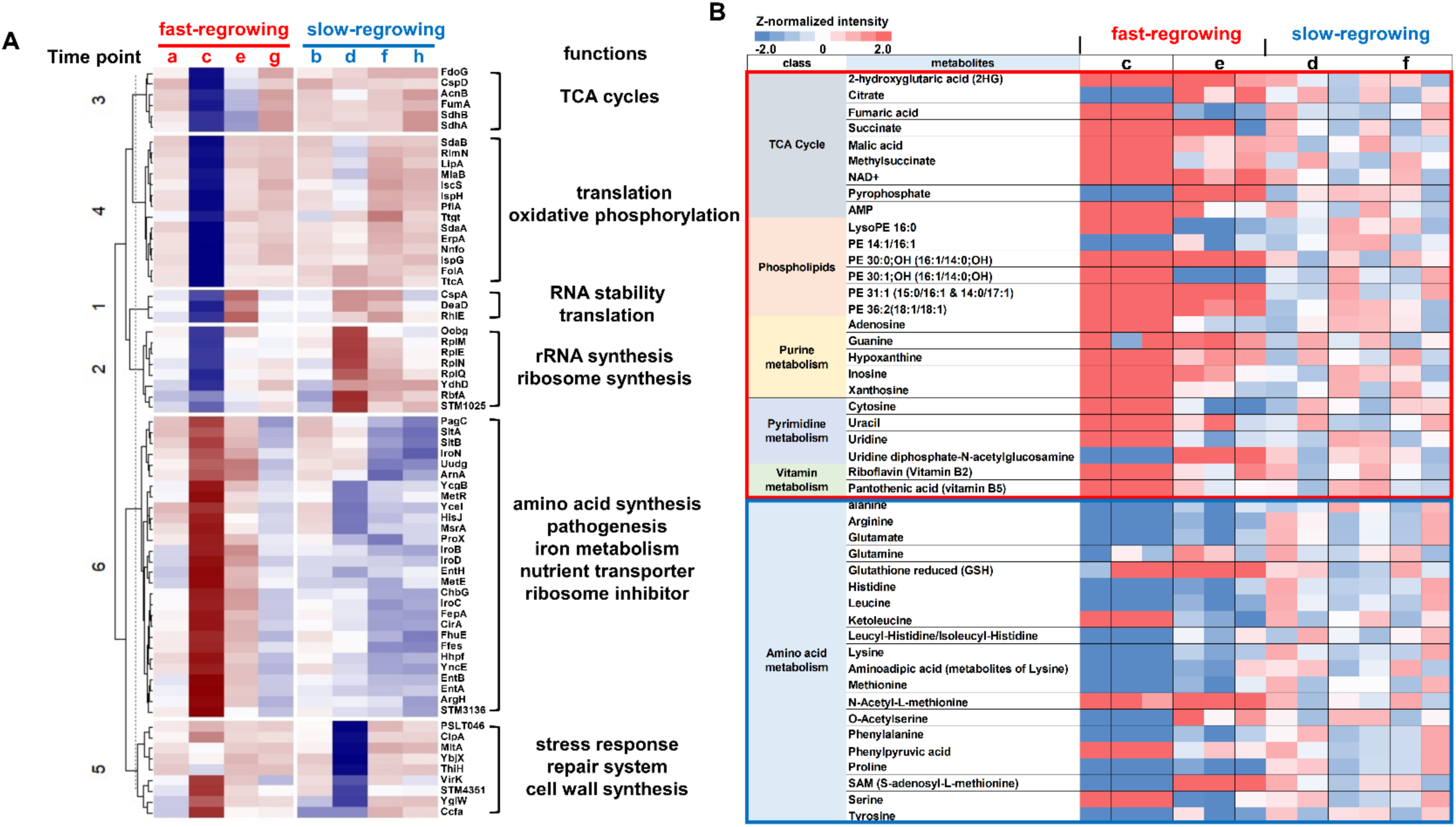
Proteomic and metabolomic profiles during dormancy in *Salmonella* Typhimurium. (A) Hierarchical clustering analysis (HCA) of proteins differentially expressed during dormancy. Time points represent dormant bacteria (a-d) and reinoculated bacteria (e-h) under fast-regrowth (red) and slow-regrowth (blue) conditions. Functional groups are annotated and include respiration, energy metabolism, RNA stability, translation, amino acid biosynthesis, stress response, and cell wall synthesis. Z-scores represent normalized protein intensities across time points. (B) Heatmap of metabolite abundance across dormant (c and d) and reinoculated (e and f) *Salmonella* Typhimurium under fast– (c and e) and slow-regrowing (d and f) dormant bacteria. Metabolites are grouped by class, including the TCA cycle, phospholipids, nucleotide metabolism, vitamins, and amino acids. Colors represent Z-score–normalized relative abundance across conditions.

**Fig. S2.**
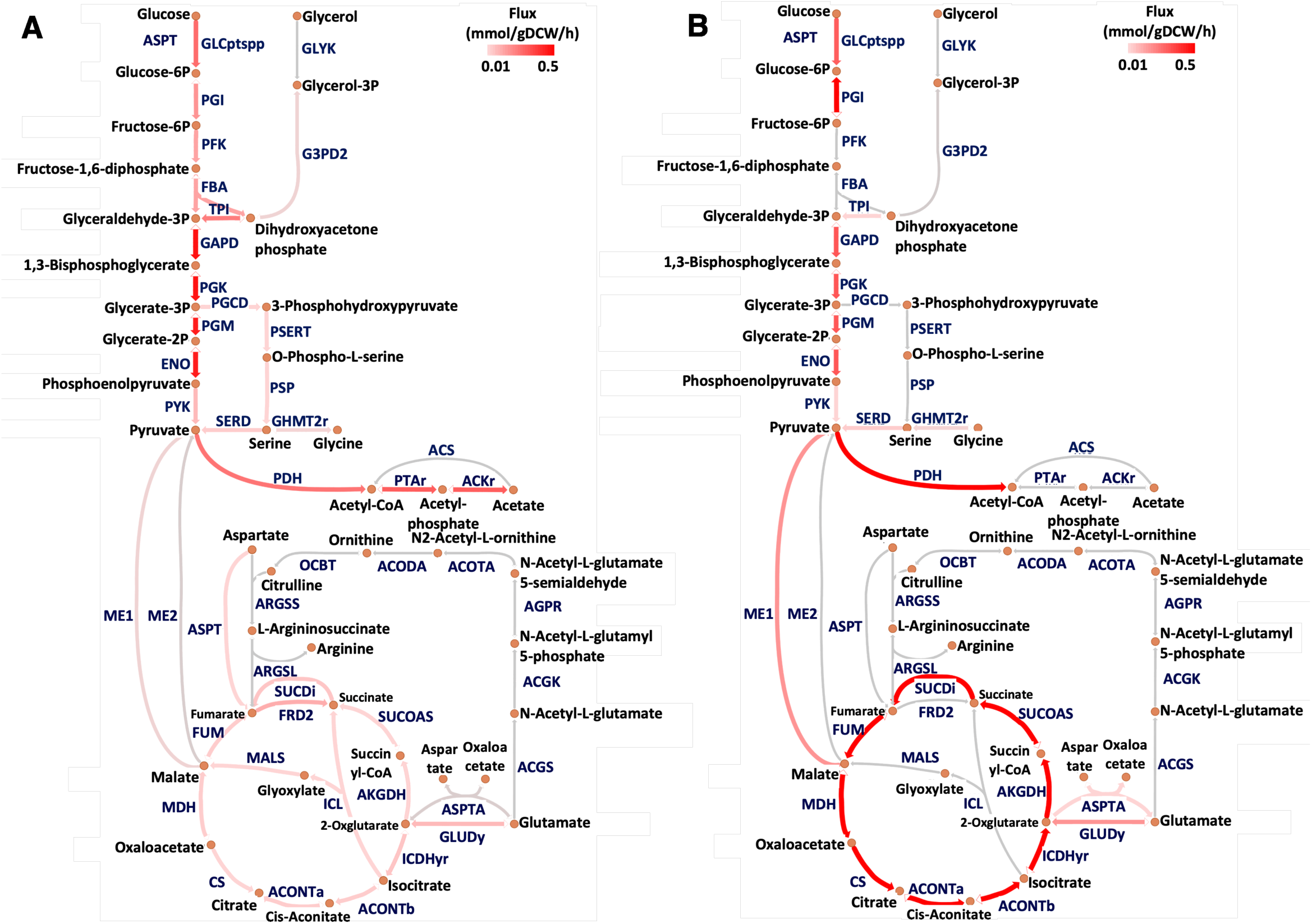
*In silico* metabolic flux analysis between fast and slow-regrowing dormant bacteria. Metabolic flux analysis was compared between fast-regrowing dormant bacteria (low-ATP) (A) and slow-regrowing dormant bacteria (high-ATP) (B) groups within central metabolic pathways, which include glycolysis, TCA cycle, serine, and arginine biosynthesis. Low ATP levels promote rapid resuscitation of dormant bacteria(*6*). Reactions are depicted as arrows with abbreviation labels in blue, and metabolites are shown as orange circles. Reactions carrying no flux are shown in grey, and both the color intensity and arrow width indicate the flux magnitude in mmol/gDCW/h. Abbreviation for metabolic enzyme. ACGK, Acetylglutamate kinase; ACGS, N-acetylglutamate synthase; ACKr, Acetate kinase; ACODA, Acetylornithine deacetylase; ACONTa, Aconitase; ACONTb, Aconitase; ACOTA, Acetylornithine transaminase; ACS, Acetyl-CoA synthetase; AGPR, N-acetyl-g-glutamyl-phosphate reductase; AGRSL, Argininosuccinate lyase; AKGDH, 2-Oxogluterate dehydrogenase; ARGSS, Argininosuccinate synthase; ASPT, L-aspartase; ASPTA, Aspartate transaminase; CS, Citrate synthase; ENO, Enolase; FBA, Fructose-bisphosphate aldolase; FRD2, Fumarate reductase; FUM, Fumarase; G3PD2, Glycerol-3-phosphate dehydrogenase; GAPD, Glyceraldehyde-3-phosphate dehydrogenase; GHMT2r, Glycine hydroxymethyltransferase reversible; GLCptspp, D-glucose transport via PEP:Pyr PTS; GLUDy, Glutamate dehydrogenase; GLYK, Glycerol kinase; ICDHyr, Isocitrate dehydrogenase; MDH, Malate dehydrogenase; ME1, Malic enzyme; ME2, Malic enzyme; OCBT, Ornithine carbamoyltransferase; PDH, Pyruvate dehydrogenase; PFK, Phosphofructokinase; PGCD, Phosphoglycerate dehydrogenase; PGI, Glucose-6-phosphate isomerase; PGK, Phosphoglycerate kinase; PGM, Phosphoglycerate mutase; PSERT, Phosphoserine transaminase; PSP, Phosphoserine phosphatase; PTAr, Phosphotransacetylase; PYK, Pyruvate kinase; SERD, L-serine deaminase; SUCDi, Succinate dehydrogenase; SUCOAS, Succinyl-CoA synthetase; TPI, Triose-phosphate isomerase.

**Fig. S3.**
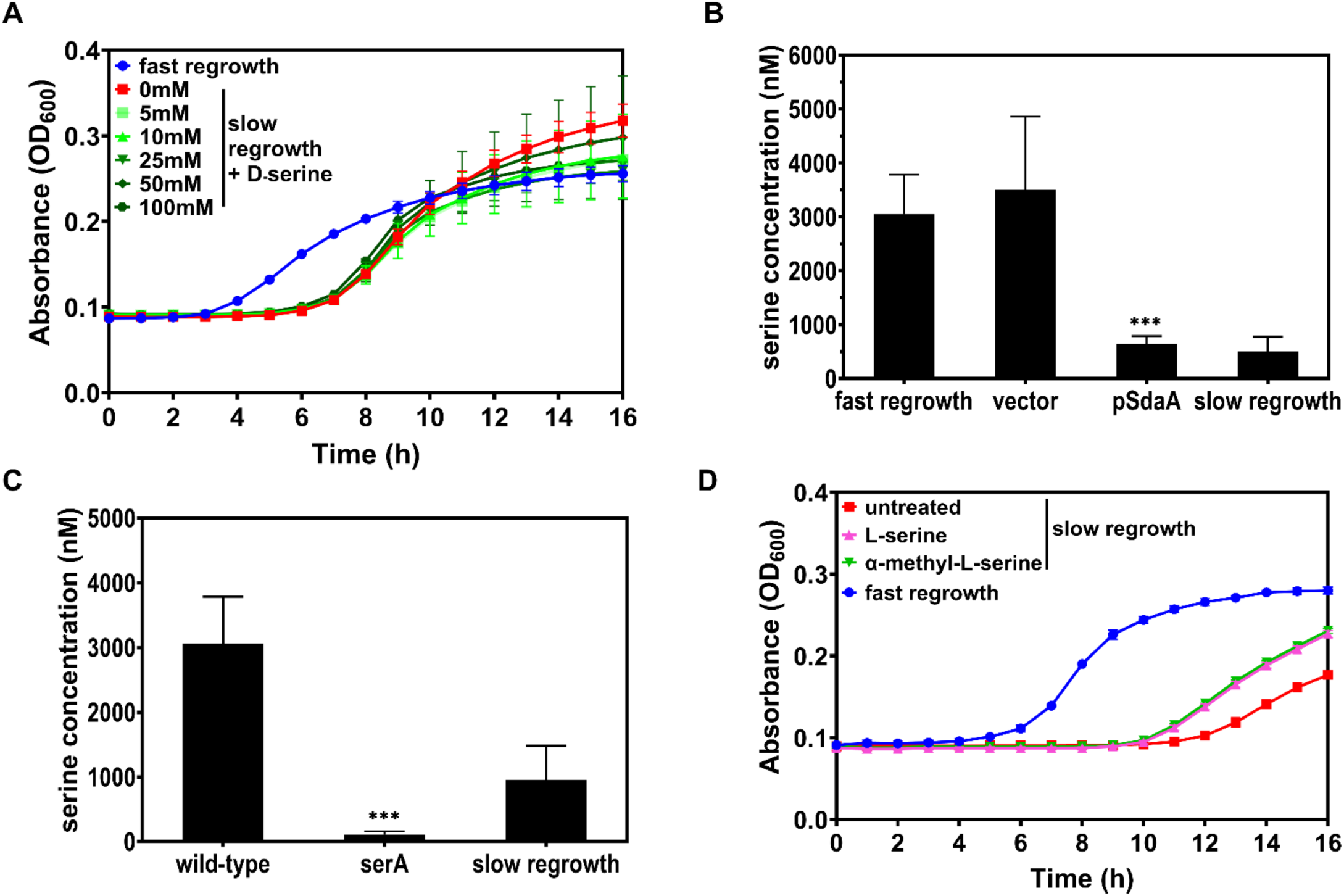
L-serine and L-serine analogs promote the regrowth of dormant bacteria. (A) Growth curves show that D-serine supplementation does not accelerate the regrowth of dormant bacteria. (B) Intracellular serine concentration is reduced in bacteria harboring a plasmid with the *sdaA* gene, which encodes a serine degradation enzyme. Unpaired Student’s t tests were performed between pSdaA samples with fast regrowth samples; *\*\*\*P* < 0.001. (C). Inactivation of *serA*, which encodes a serine biosynthesis enzyme, markedly decreases intracellular L-serine levels. Unpaired Student’s t tests were performed between *serA* samples with other combinations; *\*\*\*P* < 0.001. (D) L-serine analog alpha-methyl-L-serine promotes rapid resuscitation in dormant bacteria, as does L-serine. Growth curves indicate that supplementation of L-serine or alpha-methyl-L-serine accelerates the regrowth of dormant bacteria. The mean and SD from three independent experiments are shown.

**Fig. S4.**
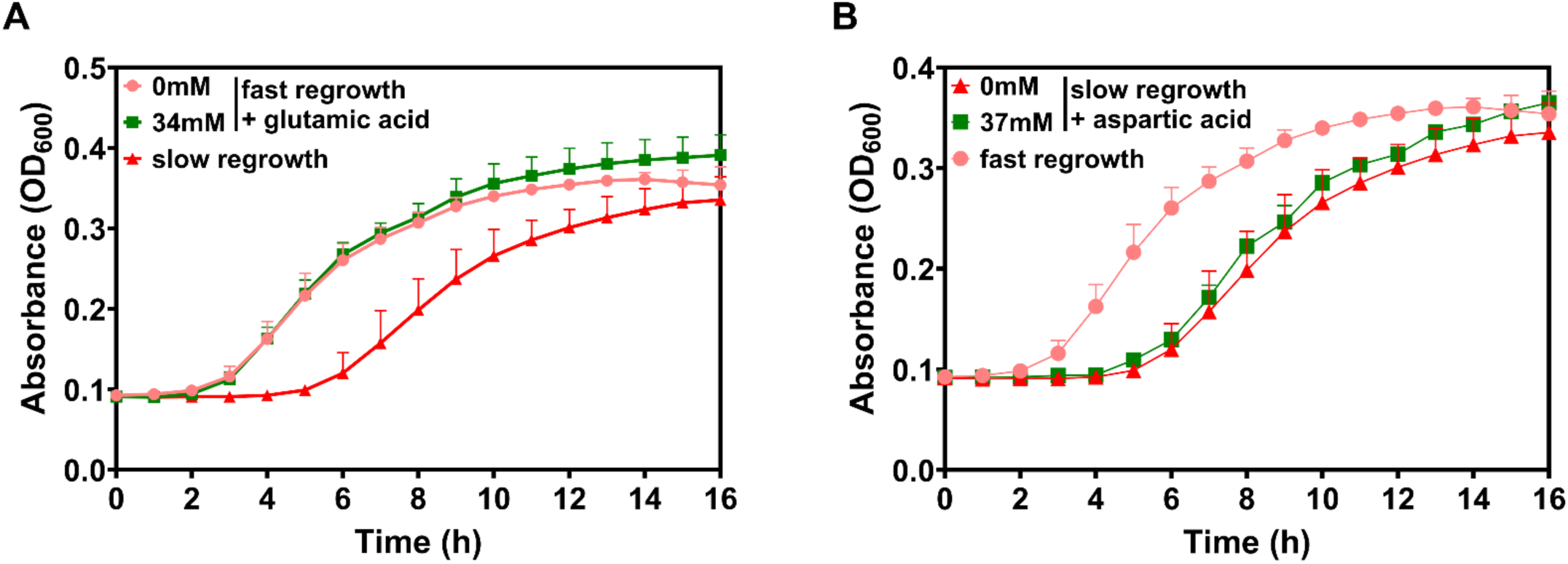
Glutamic acid or aspartic acid supplementation does not promote rapid resuscitation from dormancy. Growth curves indicate that supplementation with glutamic acid (A) or aspartic acid (B) does not accelerate regrowth of dormant bacteria.

**Fig. S5.**
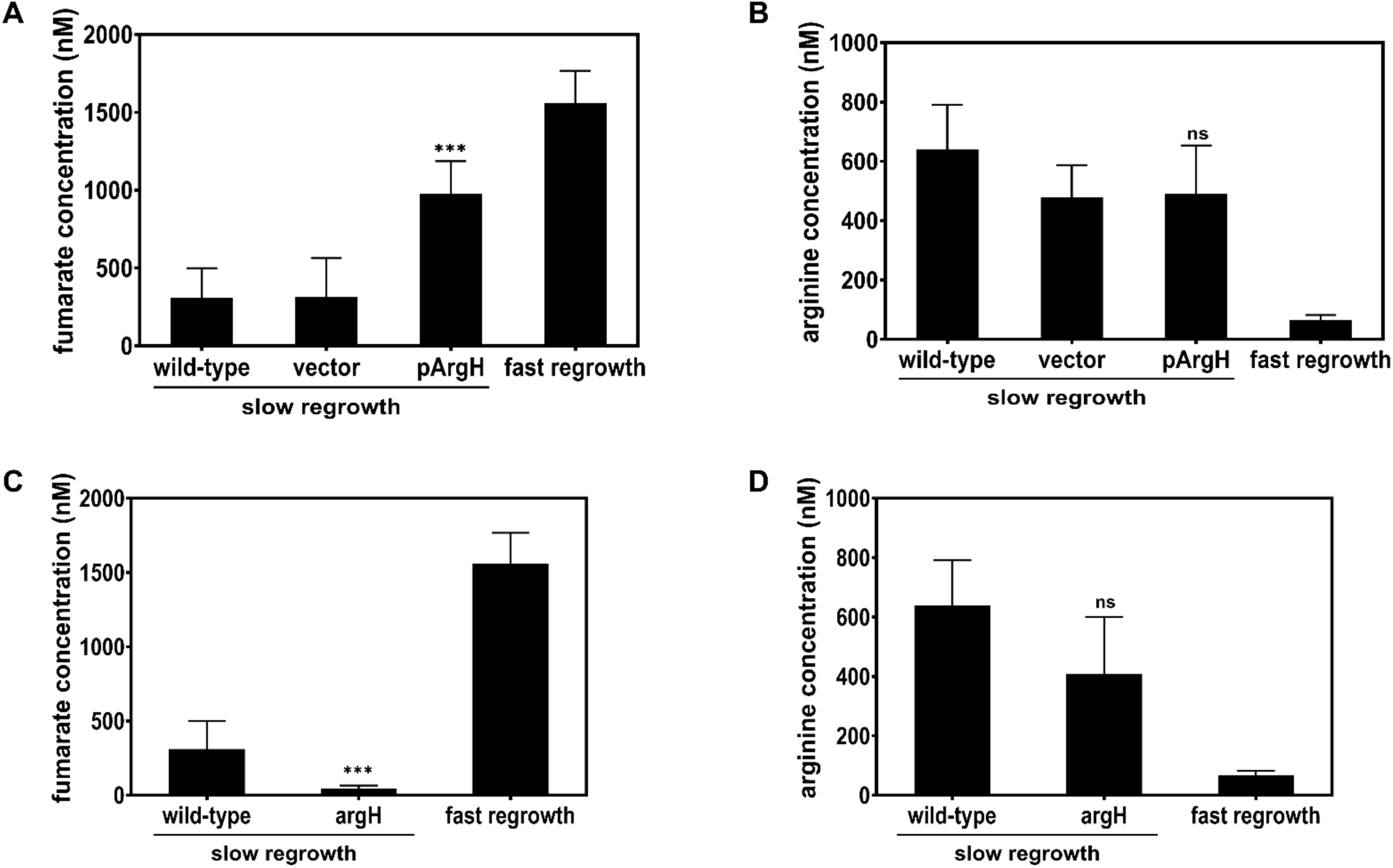
ArgH regulates the levels of arginine and fumarate during dormancy. (A and B) Intracellular concentrations of fumarate (A) and arginine (B) were measured in dormant bacteria harboring a plasmid with the *argH* gene. Unpaired Student’s t tests were performed between pArgH samples with wild-type (slow regrowth sample); *\*\*\*P* < 0.001 and ns; not significant. (C and D) Intracellular concentrations of fumarate (C) and arginine (D) were measured in bacteria with an inactivated *argH* gene during dormancy. Unpaired Student’s t tests were performed between *argH* samples with wild-type (slow regrowth sample); *\*\*\*P* < 0.001 and ns; not significant. The mean and SD from three independent experiments are shown.

**Fig. S6.**
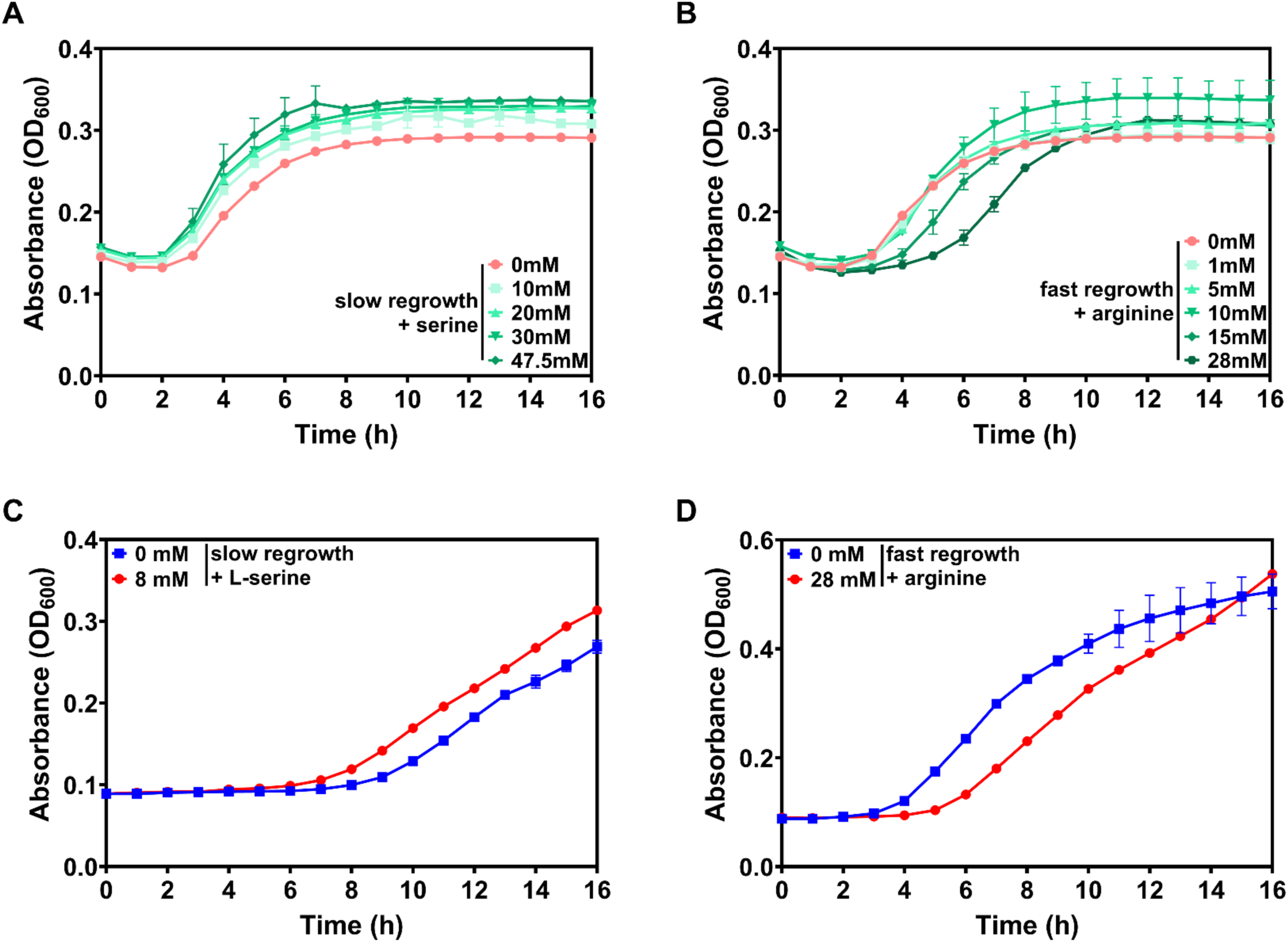
L-serine and arginine regulate the resuscitation of meropenem-induced persister *E. coli and* chloramphenicol induced dormant *methicillin resistant Staphylococcus aureus* (MRSA). (A) Growth curves indicate that supplementation with L-serine accelerates regrowth of persister *E. coli*. (B) Supplementation with arginine does not accelerate regrowth of persister *E. coli*. (C) L-serine accelerates regrowth of chloramphenicol-induced persister MRSA. (D) Supplementation with arginine prolongs dormancy of MRSA.

**Fig. S7.**
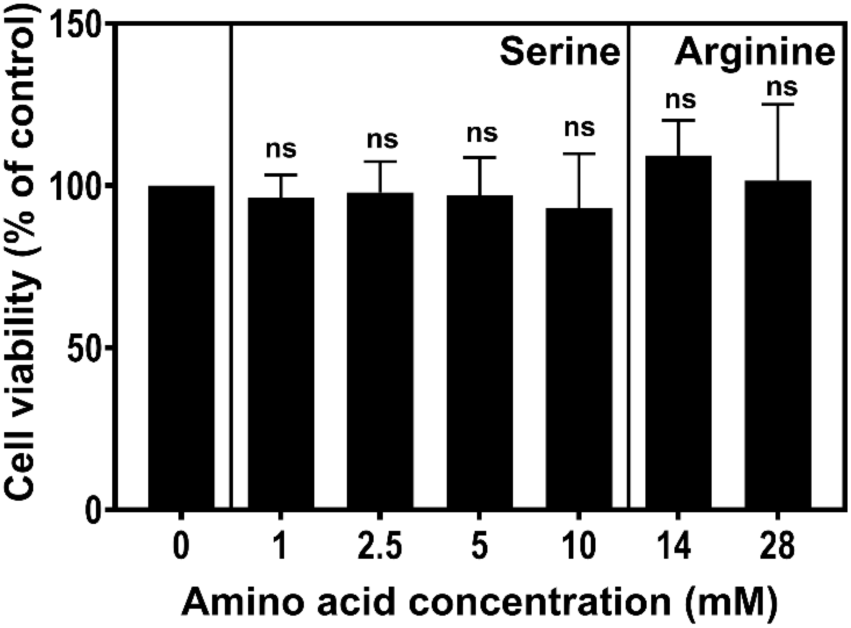
L-serine and arginine exhibit no cytotoxicity toward macrophages at experimental concentrations. J774A.1 macrophages were treated with PBS (0 mM), L-serine, or arginine at the indicated concentrations. Neither L-serine nor arginine significantly affected macrophage viability compared to the PBS samples (0 mM). Unpaired Student’s t tests were performed between each treatment group and the PBS; ns, not significant. The mean and SD from three independent experiments are shown.

**Fig. S8.**
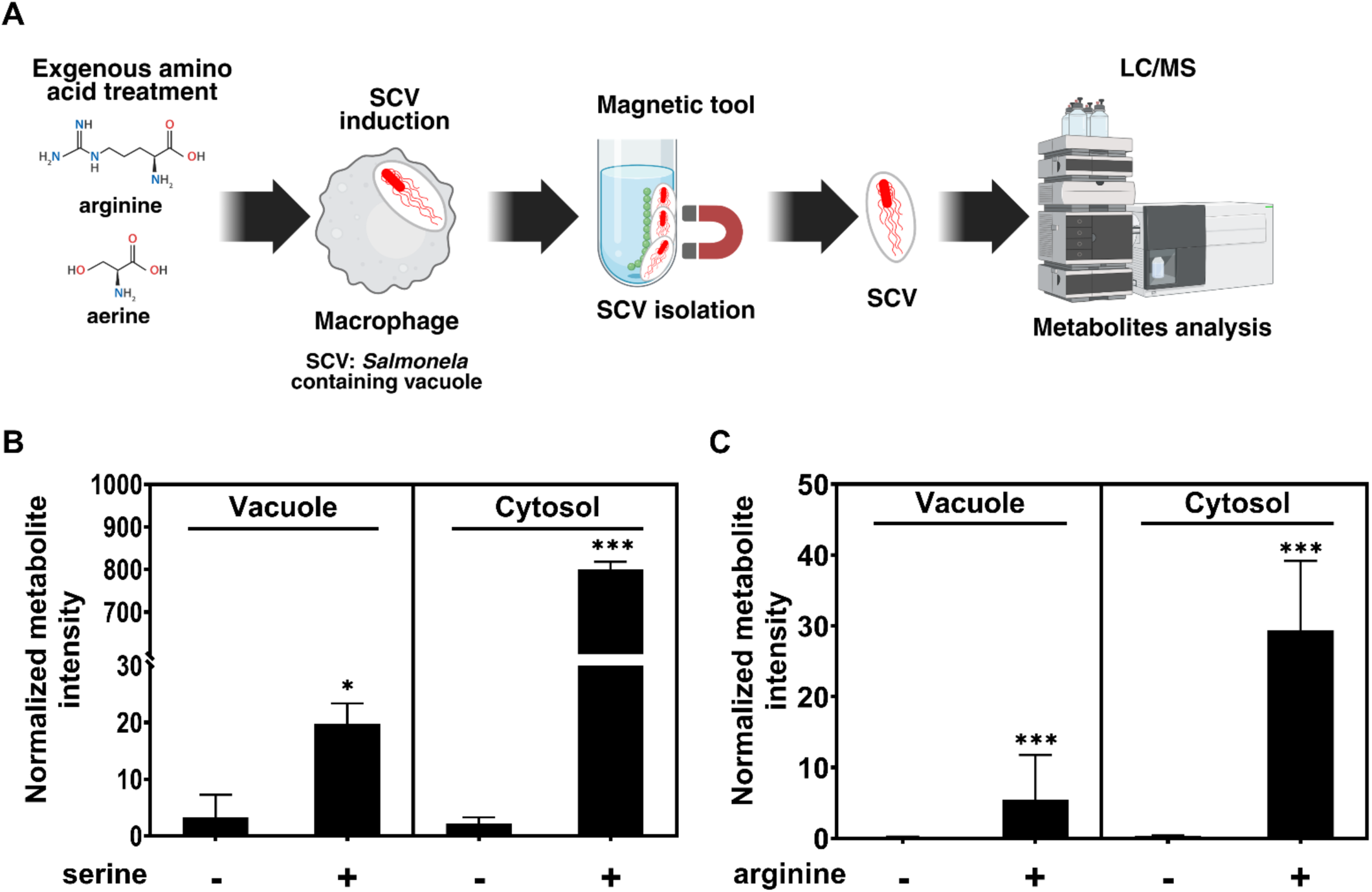
Exogenous metabolites reach the *Salmonella*-containing vacuole and modulate intracellular bacterial resuscitation. (A) Schematic illustration shows magnetic-mediated isolation of *Salmonella*-containing vacuoles (SCVs) from infected macrophages following exogenous amino acid treatment for metabolomic profiling. (B and C) Liquid chromatography-mass spectrophotometry (LC-MS) quantification of exogenous L-serine (B) and arginine (C) in SCV vacuole and cytosolic fractions, indicates the levels of metabolites in cytosol and vacuole in dormant bacteria. Unpaired Student’s t tests were performed between amino acid-treated (+) and untreated (−) groups within each fraction; \**P* < 0.05, \*\*\**P* < 0.001. The mean and SD from three independent experiments are shown.

## Supplemental Tables

**Table S1.** Microbial strains and plasmids used in this study.

| Strains | Relevant characteristics | Source |
| --- | --- | --- |
| *Escherichia coli* |  |  |
| DH5a | Host strain used for generation and propagation of plasmid construct | Addgene |
| HN100 | K-12 MG1655 wild-type | (76) |
| *Salmonella enterica* serovar Typhimurium |  |  |
| 14028s | wild-type | (77) |
| TY13 | pUHE-21-2-*lacI*<sup>q</sup> | This study |
| TY44 | *sdaA*-HA::km | This study |
| TY45 | *serA*-HA::km | This study |
| TY40 | *serA* | This study |
| TY23 | pUHE-21-2-*lacI*<sup>q</sup>::*sdaA* | This study |
| TY46 | *argH*-HA::km | This study |
| TY32 | *argH*::km | This study |
| TY37 | *argH* | This study |
| TY22 | pUHE-21-2-*lacI*<sup>q</sup>::*argH* | This study |
| *Staphylococcus aureus* |  |  |
| JE25 | Methicillin-resistant *Staphylococcus aureus* | Seoul National University Bundang Hospital |
| **Plasmids** |  |  |
| pKD46 | rep<sub>pSC101</sub><sup>ts</sup> Amp<sup>R</sup> P<sub>araBAD</sub>-gbexo | (78) |
| pKD4 | rep<sub>R6Kg</sub> Amp<sup>R</sup> FRT Km<sup>R</sup> FRT | (78) |
| pCP20 | rep<sub>pSC101</sub><sup>ts</sup> 1 cI857 FLP Amp<sup>R</sup> Cm<sup>R</sup> | (78) |
| pUHE-21-2-*lacI*<sup>q</sup> | Rep<sub>pMB1</sub> *lacI*<sup>q</sup> Amp<sup>R</sup>, pATPase vector control | (79) |

**Table S2.**
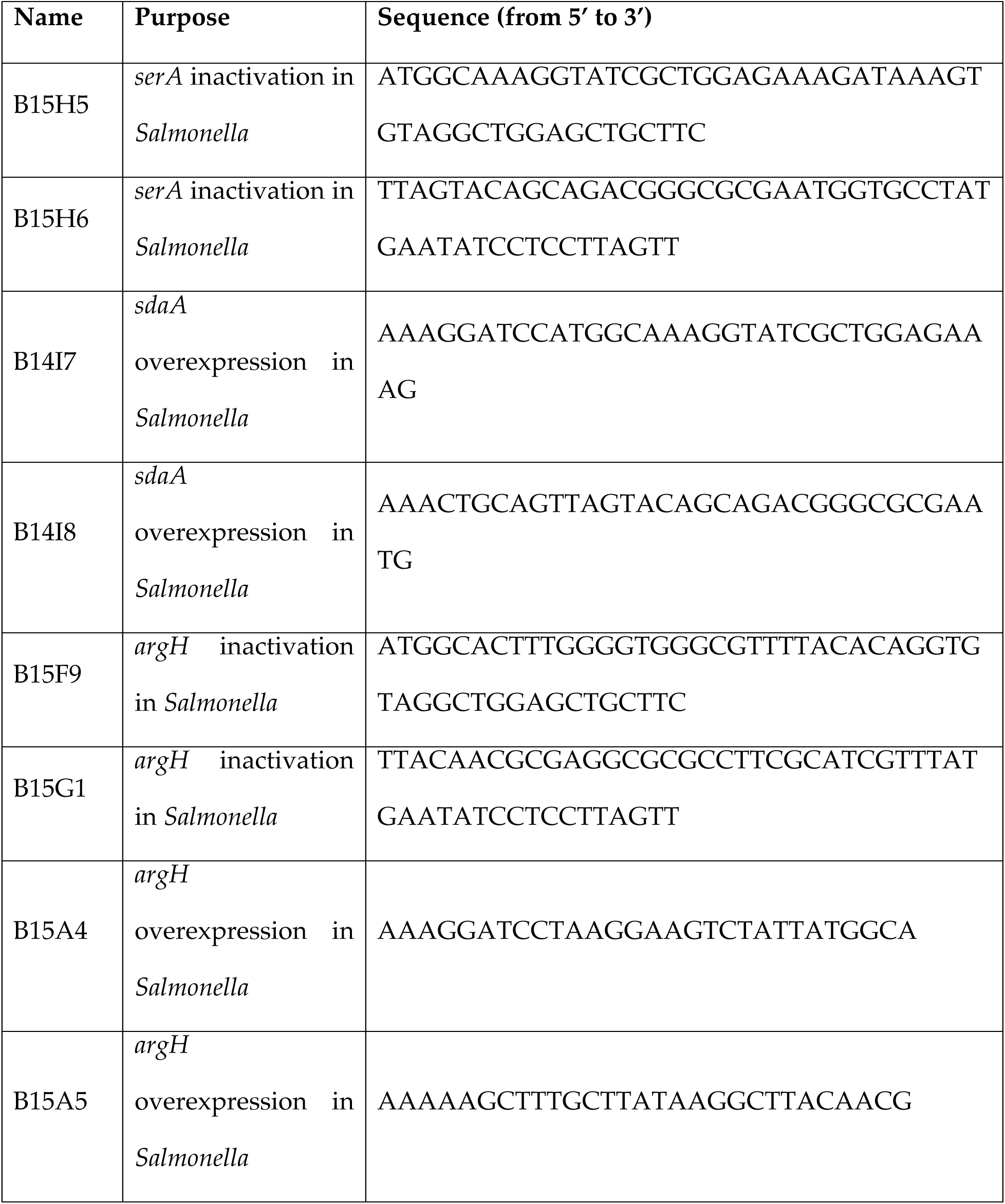
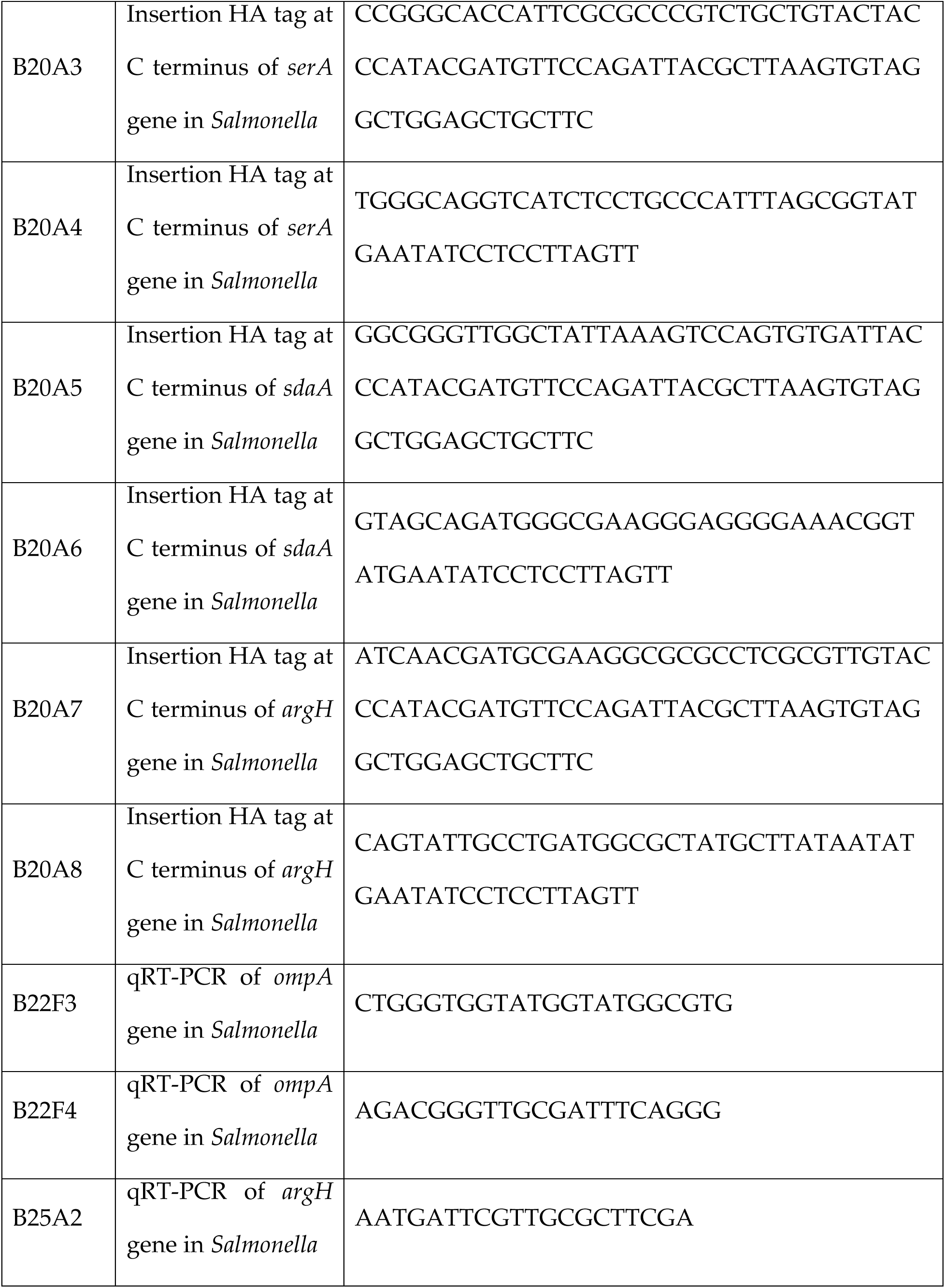
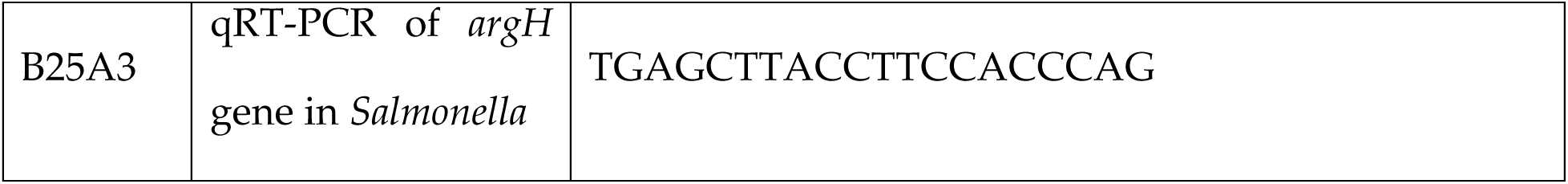
Primers used for strain construction and quantitative RT-PCR.

## References

1. N. Aviram, A. K. Shilton, N. G. Lyn, B. S. Reis, A. Brivanlou, L. A. Marraffini, Cas10 relieves host growth arrest to facilitate spacer retention during type III-A CRISPR-Cas immunity. Cell Host Microbe 32, 2050–2062 e2056 (2024).

2. J. Fares, M. Y. Fares, H. H. Khachfe, H. A. Salhab, Y. Fares, Molecular principles of metastasis: a hallmark of cancer revisited. Signal Transduct Target Ther 5, 28 (2020).

3. R. Kolter, N. Balaban, T. Julou, Bacteria grow swiftly and live thriftily. Curr Biol 32, R599–R605 (2022).

4. E. Kussell, R. Kishony, N. Q. Balaban, S. Leibler, Bacterial persistence: a model of survival in changing environments. Genetics 169, 1807–1814 (2005).

5. S. Wagley, H. Morcrette, A. Kovacs-Simon, Z. R. Yang, A. Power, R. K. Tennant, J. Love, N. Murray, R. W. Titball, C. S. Butler, Bacterial dormancy: A subpopulation of viable but non-culturable cells demonstrates better fitness for revival. PLoS Pathog 17, e1009194 (2021).

6. J. Yeom, E. A. Groisman, Reduced ATP-dependent proteolysis of functional proteins during nutrient limitation speeds the return of microbes to a growth state. Sci Signal 14, (2021).

7. V. Chubukov, L. Gerosa, K. Kochanowski, U. Sauer, Coordination of microbial metabolism. Nat Rev Microbiol 12, 327–340 (2014).

8. M. Bonora, S. Patergnani, A. Rimessi, E. De Marchi, J. M. Suski, A. Bononi, C. Giorgi, S. Marchi, S. Missiroli, F. Poletti, M. R. Wieckowski, P. Pinton, ATP synthesis and storage. Purinergic Signal 8, 343–357 (2012).

9. E. Noor, E. Eden, R. Milo, U. Alon, Central carbon metabolism as a minimal biochemical walk between precursors for biomass and energy. Mol Cell 39, 809–820 (2010).

10. F. Rojo, Carbon catabolite repression in Pseudomonas: optimizing metabolic versatility and interactions with the environment. FEMS Microbiol Rev 34, 658–684 (2010).

11. L. K. Do, H. M. Lee, Y. S. Ha, C. H. Lee, J. Kim, Amino acids in cancer: Understanding metabolic plasticity and divergence for better therapeutic approaches. Cell Rep 44, 115529 (2025).

12. N. S. Chandel, Amino Acid Metabolism. Cold Spring Harb Perspect Biol 13, (2021).

13. Y. Zhang, Z. Lin, Q. Liu, Y. Li, Z. Wang, H. Ma, T. Chen, X. Zhao, Engineering of Serine-Deamination pathway, Entner-Doudoroff pathway and pyruvate dehydrogenase complex to improve poly(3-hydroxybutyrate) production in Escherichia coli. Microb Cell Fact 13, 172 (2014).

14. M. Papagianni, Recent advances in engineering the central carbon metabolism of industrially important bacteria. Microb Cell Fact 11, 50 (2012).

15. W. Eisenreich, T. Dandekar, J. Heesemann, W. Goebel, Carbon metabolism of intracellular bacterial pathogens and possible links to virulence. Nat Rev Microbiol 8, 401–412 (2010).

16. L. Rohmer, D. Hocquet, S. I. Miller, Are pathogenic bacteria just looking for food? Metabolism and microbial pathogenesis. Trends Microbiol 19, 341–348 (2011).

17. R. A. Fisher, B. Gollan, S. Helaine, Persistent bacterial infections and persister cells. Nat Rev Microbiol 15, 453–464 (2017).

18. B. Van den Bergh, J. E. Michiels, T. Wenseleers, E. M. Windels, P. V. Boer, D. Kestemont, L. De Meester, K. J. Verstrepen, N. Verstraeten, M. Fauvart, J. Michiels, Frequency of antibiotic application drives rapid evolutionary adaptation of Escherichia coli persistence. Nat Microbiol 1, 16020 (2016).

19. P. W. S. Hill, A. L. Moldoveanu, M. Sargen, S. Ronneau, I. Glegola-Madejska, C. Beetham, R. A. Fisher, S. Helaine, The vulnerable versatility of Salmonella antibiotic persisters during infection. Cell Host Microbe 29, 1757–1773 e1710 (2021).

20. N. Q. Balaban, J. Merrin, R. Chait, L. Kowalik, S. Leibler, Bacterial persistence as a phenotypic switch. Science 305, 1622–1625 (2004).

21. H. Niu, J. Gu, Y. Zhang, Bacterial persisters: molecular mechanisms and therapeutic development. Signal Transduct Target Ther 9, 174 (2024).

22. B. P. Conlon, S. E. Rowe, A. B. Gandt, A. S. Nuxoll, N. P. Donegan, E. A. Zalis, G. Clair, J. N. Adkins, A. L. Cheung, K. Lewis, Persister formation in Staphylococcus aureus is associated with ATP depletion. Nat Microbiol 1, (2016).

23. Y. Pu, Y. Li, X. Jin, T. Tian, Q. Ma, Z. Zhao, S. Y. Lin, Z. Chen, B. Li, G. Yao, M. C. Leake, C. J. Lo, F. Bai, ATP-Dependent Dynamic Protein Aggregation Regulates Bacterial Dormancy Depth Critical for Antibiotic Tolerance. Mol Cell 73, 143–156 e144 (2019).

24. J. Wainwright, G. Hobbs, I. Nakouti, Persister cells: formation, resuscitation and combative therapies. Arch Microbiol 203, 5899–5906 (2021).

25. J. Qi, X. Xiao, L. Ouyang, C. Yang, Y. Zhuang, L. Zhang, Enhancement of fatty acid degradation pathway promoted glucoamylase synthesis in Aspergillus niger. Microb Cell Fact 21, 238 (2022).

26. L. Artzi, A. Alon, K. P. Brock, A. G. Green, A. Tam, F. H. Ramirez-Guadiana, D. Marks, A. Kruse, D. Z. Rudner, Dormant spores sense amino acids through the B subunits of their germination receptors. Nat Commun 12, 6842 (2021).

27. H. Nozaki, S. Kuroda, K. Watanabe, K. Yokozeki, Purification and gene cloning of alpha-methylserine aldolase from Ralstonia sp. strain AJ110405 and application of the enzyme in the synthesis of alpha-methyl-L-serine. Appl Environ Microbiol 74, 7596–7599 (2008).

28. G. Kikuchi, Y. Motokawa, T. Yoshida, K. Hiraga, Glycine cleavage system: reaction mechanism, physiological significance, and hyperglycinemia. Proc Jpn Acad Ser B Phys Biol Sci 84, 246–263 (2008).

29. I. Sanchez-Andrea, I. A. Guedes, B. Hornung, S. Boeren, C. E. Lawson, D. Z. Sousa, A. Bar-Even, N. J. Claassens, A. J. M. Stams, The reductive glycine pathway allows autotrophic growth of Desulfovibrio desulfuricans. Nat Commun 11, 5090 (2020).

30. B. Moosavi, E. A. Berry, X. L. Zhu, W. C. Yang, G. F. Yang, The assembly of succinate dehydrogenase: a key enzyme in bioenergetics. Cell Mol Life Sci 76, 4023–4042 (2019).

31. H. Shirai, K. Mizuguchi, Prediction of the structure and function of AstA and AstB, the first two enzymes of the arginine succinyltransferase pathway of arginine catabolism. FEBS Lett 555, 505–510 (2003).

32. S. Wuttge, M. Bommer, F. Jager, B. M. Martins, S. Jacob, A. Licht, F. Scheffel, H. Dobbek, E. Schneider, Determinants of substrate specificity and biochemical properties of the sn-glycerol-3-phosphate ATP binding cassette transporter (UgpB-AEC2) of Escherichia coli. Mol Microbiol 86, 908–920 (2012).

33. A. Tikhomirova, M. M. Rahman, S. P. Kidd, R. L. Ferrero, A. Roujeinikova, Cysteine and resistance to oxidative stress: implications for virulence and antibiotic resistance. Trends Microbiol 32, 93–104 (2024).

34. J. Ramoneda, K. Fan, J. M. Lucas, H. Chu, A. Bissett, M. S. Strickland, N. Fierer, Ecological relevance of flagellar motility in soil bacterial communities. ISME J 18, (2024).

35. C. Cheng, Z. Dong, X. Han, J. Sun, H. Wang, L. Jiang, Y. Yang, T. Ma, Z. Chen, J. Yu, W. Fang, H. Song, Listeria monocytogenes 10403S Arginine Repressor ArgR Finely Tunes Arginine Metabolism Regulation under Acidic Conditions. Front Microbiol 8, 145 (2017).

36. A. W. Crocker, C. E. Harty, J. H. Hammond, S. D. Willger, P. Salazar, N. J. Botelho, N. J. Jacobs, D. A. Hogan, Pseudomonas aeruginosa Ethanol Oxidation by AdhA in Low-Oxygen Environments. J Bacteriol 201, (2019).

37. X. Fang, K. R. Allison, Resuscitation dynamics reveal persister partitioning after antibiotic treatment. Mol Syst Biol 19, e11320 (2023).

38. R. L. Santos, A. J. Baumler, Cell tropism of Salmonella enterica. Int J Med Microbiol 294, 225–233 (2004).

39. S. Helaine, A. M. Cheverton, K. G. Watson, L. M. Faure, S. A. Matthews, D. W. Holden, Internalization of Salmonella by macrophages induces formation of nonreplicating persisters. Science 343, 204–208 (2014).

40. B. Ekinci, A. Y. Coban, A. Birinci, B. Durupinar, M. Erturk, In vitro effects of cefotaxime and ceftriaxone on Salmonella typhi within human monocyte-derived macrophages. Clin Microbiol Infect 8, 810–813 (2002).

41. K. D. Passalacqua, M. E. Charbonneau, M. X. D. O’Riordan, Bacterial Metabolism Shapes the Host-Pathogen Interface. Microbiol Spectr 4, (2016).

42. S. Kitamoto, C. J. Alteri, M. Rodrigues, H. Nagao-Kitamoto, K. Sugihara, S. D. Himpsl, M. Bazzi, M. Miyoshi, T. Nishioka, A. Hayashi, T. L. Morhardt, P. Kuffa, H. Grasberger, M. El-Zaatari, S. Bishu, C. Ishii, A. Hirayama, K. A. Eaton, B. Dogan, K. W. Simpson, N. Inohara, H. L. T. Mobley, J. Y. Kao, S. Fukuda, N. Barnich, N. Kamada, Dietary L-serine confers a competitive fitness advantage to Enterobacteriaceae in the inflamed gut. Nat Microbiol 5, 116–125 (2020).

43. C. C. Y. Chan, I. A. Lewis, Role of metabolism in uropathogenic Escherichia coli. Trends Microbiol 30, 1174–1204 (2022).

44. M. Yasugi, A. Ohta, K. Takano, K. Yakubo, M. Irie, M. Miyake, Serine affects engulfment during the sporulation process in Clostridium perfringens strain SM101. Anaerobe 90, 102914 (2024).

45. X. Zhou, L. He, C. Wu, Y. Zhang, X. Wu, Y. Yin, Serine alleviates oxidative stress via supporting glutathione synthesis and methionine cycle in mice. Mol Nutr Food Res 61, (2017).

46. D. X. Yang, M. J. Yang, Y. Yin, T. S. Kou, L. T. Peng, Z. G. Chen, J. Zheng, B. Peng, Serine Metabolism Tunes Immune Responses To Promote Oreochromis niloticus Survival upon Edwardsiella tarda Infection. mSystems 6, e0042621 (2021).

47. B. L. Schneider, A. K. Kiupakis, L. J. Reitzer, Arginine catabolism and the arginine succinyltransferase pathway in Escherichia coli. J Bacteriol 180, 4278–4286 (1998).

48. T. Rimaux, A. Riviere, K. Illeghems, S. Weckx, L. De Vuyst, F. Leroy, Expression of the arginine deiminase pathway genes in Lactobacillus sakei is strain dependent and is affected by the environmental pH. Appl Environ Microbiol 78, 4874–4883 (2012).

49. A. P. Snell, D. A. Manias, R. R. Elbehery, G. M. Dunny, J. L. E. Willett, Arginine impacts aggregation, biofilm formation, and antibiotic susceptibility in Enterococcus faecalis. FEMS Microbes 5, xtae030 (2024).

50. Y. Liu, S. Liu, Q. Zhi, P. Zhuang, R. Zhang, Z. Zhang, K. Zhang, Y. Sun, Arginine-induced metabolomic perturbation in Streptococcus mutans. J Oral Microbiol 14, 2015166 (2022).

51. H. Van Acker, P. Van Dijck, T. Coenye, Molecular mechanisms of antimicrobial tolerance and resistance in bacterial and fungal biofilms. Trends Microbiol 22, 326–333 (2014).

52. P. S. Stewart, M. J. Franklin, Physiological heterogeneity in biofilms. Nat Rev Microbiol 6, 199–210 (2008).

53. J. J. Lemke, P. Sanchez-Vazquez, H. L. Burgos, G. Hedberg, W. Ross, R. L. Gourse, Direct regulation of Escherichia coli ribosomal protein promoters by the transcription factors ppGpp and DksA. Proc Natl Acad Sci U S A 108, 5712–5717 (2011).

54. M. Zhu, H. Mu, X. Dai, Integrated control of bacterial growth and stress response by (p)ppGpp in Escherichia coli: A seesaw fashion. iScience 27, 108818 (2024).

55. A. Lichev, A. Angelov, I. Cucurull, W. Liebl, Amino acids as nutritional factors and (p)ppGpp as an alarmone of the stringent response regulate natural transformation in Micrococcus luteus. Sci Rep 9, 11030 (2019).

56. C. S. Schaumburg, M. Tan, Arginine-dependent gene regulation via the ArgR repressor is species specific in chlamydia. J Bacteriol 188, 919–927 (2006).

57. M. Huemer, S. Mairpady Shambat, S. D. Brugger, A. S. Zinkernagel, Antibiotic resistance and persistence-Implications for human health and treatment perspectives. EMBO Rep 21, e51034 (2020).

58. A. L. Moldoveanu, J. A. Rycroft, S. Helaine, Impact of bacterial persisters on their host. Curr Opin Microbiol 59, 65–71 (2021).

59. N. Q. Balaban, S. Helaine, K. Lewis, M. Ackermann, B. Aldridge, D. I. Andersson, M. P. Brynildsen, D. Bumann, A. Camilli, J. J. Collins, C. Dehio, S. Fortune, J. M. Ghigo, W. D. Hardt, A. Harms, M. Heinemann, D. T. Hung, U. Jenal, B. R. Levin, J. Michiels, G. Storz, M. W. Tan, T. Tenson, L. Van Melderen, A. Zinkernagel, Definitions and guidelines for research on antibiotic persistence. Nat Rev Microbiol 17, 441–448 (2019).

60. B. Peng, Y. B. Su, H. Li, Y. Han, C. Guo, Y. M. Tian, X. X. Peng, Exogenous alanine and/or glucose plus kanamycin kills antibiotic-resistant bacteria. Cell Metab 21, 249–262 (2015).

61. S. Ronneau, C. Michaux, S. Helaine, Decline in nitrosative stress drives antibiotic persister regrowth during infection. Cell Host Microbe 31, 993–1006 e1006 (2023).

62. P. Bhargava, J. J. Collins, Boosting bacterial metabolism to combat antibiotic resistance. Cell Metab 21, 154–155 (2015).

63. S. Tau, T. W. Miller, The role of cancer cell bioenergetics in dormancy and drug resistance. Cancer Metastasis Rev 42, 87–98 (2023).

64. Q. Lin, H. T. Tan, M. C. M. Chung, Next Generation Proteomics for Clinical Biomarker Detection Using SWATH-MS. Methods Mol Biol 1977, 3–15 (2019).

65. D. Han, J. Jin, J. Woo, H. Min, Y. Kim, Proteomic analysis of mouse astrocytes and their secretome by a combination of FASP and StageTip-based, high pH, reversed-phase fractionation. Proteomics 14, 1604–1609 (2014).

66. D. Han, S. Moon, Y. Kim, J. Kim, J. Jin, Y. Kim, In-depth proteomic analysis of mouse microglia using a combination of FASP and StageTip-based, high pH, reversed-phase fractionation. Proteomics 13, 2984–2988 (2013).

67. J. R. Wisniewski, F. Z. Gaugaz, Fast and sensitive total protein and Peptide assays for proteomic analysis. Anal Chem 87, 4110–4116 (2015).

68. S. I. Kim, H. Kim, K. Dan, H. B. Park, C. Lee, H. S. Kim, H. H. Chung, J. W. Kim, N. H. Park, D. Han, M. Lee, Proteomic landscaping of high-grade serous ovarian carcinoma identifies stearoyl-CoA desaturase 5 as a potential predictive biomarker for poly(ADP-ribose) polymerase inhibitor response. Clin Transl Med 14, e1693 (2024).

69. A. T. Kong, F. V. Leprevost, D. M. Avtonomov, D. Mellacheruvu, A. I. Nesvizhskii, MSFragger: ultrafast and comprehensive peptide identification in mass spectrometry-based proteomics. Nat Methods 14, 513–520 (2017).

70. S. Tyanova, T. Temu, P. Sinitcyn, A. Carlson, M. Y. Hein, T. Geiger, M. Mann, J. Cox, The Perseus computational platform for comprehensive analysis of (prote)omics data. Nat Methods 13, 731–740 (2016).

71. S. Tian, C. Wang, L. Yang, Y. Zhang, T. Tang, Comparison of Five Extraction Methods for Intracellular Metabolites of Salmonella typhimurium. Curr Microbiol 76, 1247–1255 (2019).

72. L. Heirendt, S. Arreckx, T. Pfau, S. N. Mendoza, A. Richelle, A. Heinken, H. S. Haraldsdottir, J. Wachowiak, S. M. Keating, V. Vlasov, S. Magnusdottir, C. Y. Ng, G. Preciat, A. Zagare, S. H. J. Chan, M. K. Aurich, C. M. Clancy, J. Modamio, J. T. Sauls, A. Noronha, A. Bordbar, B. Cousins, D. C. El Assal, L. V. Valcarcel, I. Apaolaza, S. Ghaderi, M. Ahookhosh, M. Ben Guebila, A. Kostromins, N. Sompairac, H. M. Le, D. Ma, Y. Sun, L. Wang, J. T. Yurkovich, M. A. P. Oliveira, P. T. Vuong, L. P. El Assal, I. Kuperstein, A. Zinovyev, H. S. Hinton, W. A. Bryant, F. J. Aragon Artacho, F. J. Planes, E. Stalidzans, A. Maass, S. Vempala, M. Hucka, M. A. Saunders, C. D. Maranas, N. E. Lewis, T. Sauter, B. O. Palsson, I. Thiele, R. M. T. Fleming, Creation and analysis of biochemical constraint-based models using the COBRA Toolbox v.3.0. Nat Protoc 14, 639–702 (2019).

73. L. Koduru, M. Lakshmanan, Y. Q. Lee, P. L. Ho, P. Y. Lim, W. X. Ler, S. K. Ng, D. Kim, D. S. Park, M. Banu, D. S. W. Ow, D. Y. Lee, Systematic evaluation of genome-wide metabolic landscapes in lactic acid bacteria reveals diet– and strain-specific probiotic idiosyncrasies. Cell Rep 41, 111735 (2022).

74. S. K. Kim, M. Lee, Y. Q. Lee, H. J. Lee, M. Rho, Y. Kim, J. Y. Seo, S. H. Youn, S. J. Hwang, N. G. Kang, C. H. Lee, S. Y. Park, D. Y. Lee, Genome-scale metabolic modeling and in silico analysis of opportunistic skin pathogen Cutibacterium acnes. Front Cell Infect Microbiol 13, 1099314 (2023).

75. E. Rowe, B. O. Palsson, Z. A. King, Escher-FBA: a web application for interactive flux balance analysis. BMC Syst Biol 12, 84 (2018).

76. F. R. Blattner, G. Plunkett, 3rd, C. A. Bloch, N. T. Perna, V. Burland, M. Riley, J. Collado-Vides, J. D. Glasner, C. K. Rode, G. F. Mayhew, J. Gregor, N. W. Davis, H. A. Kirkpatrick, M. A. Goeden, D. J. Rose, B. Mau, Y. Shao, The complete genome sequence of Escherichia coli K-12. Science 277, 1453–1462 (1997).

77. P. I. Fields, R. V. Swanson, C. G. Haidaris, F. Heffron, Mutants of Salmonella typhimurium that cannot survive within the macrophage are avirulent. Proc Natl Acad Sci U S A 83, 5189–5193 (1986).

78. K. A. Datsenko, B. L. Wanner, One-step inactivation of chromosomal genes in Escherichia coli K-12 using PCR products. Proc Natl Acad Sci U S A 97, 6640–6645 (2000).

79. F. C. Soncini, E. G. Vescovi, E. A. Groisman, Transcriptional autoregulation of the Salmonella typhimurium phoPQ operon. J Bacteriol 177, 4364–4371 (1995).

